# Implications of Immune-Mediated Metastatic Growth on Metastatic Dormancy, Blow-Up, Early Detection, and Treatment

**DOI:** 10.1101/814095

**Authors:** Adam Rhodes, Thomas Hillen

**Affiliations:** Department of Mathematical and Statistical Sciences, University of Alberta, Edmonton, AB, Canada

**Keywords:** Mathematical oncology, Cancer modeling, Metastasis, Dormancy, Geometric singular perturbation analysis, Ordinary differential equations

## Abstract

Metastatic seeding of distant organs can occur in the very early stages of primary tumor development. Once seeded, these micrometastases may enter a dormant phase that can last decades. Curiously, the surgical removal of the primary tumor can stimulate the accelerated growth of distant metastases, a phenomenon known as *metastatic blow-up*. Although several theories have been proposed to explain metastatic *dormancy* and *blow-up*, most mathematical investigations have ignored the important pro-tumor effects of the immune system. In this work, we address that shortcoming by developing an ordinary differential equation model of the immune-mediated theory of metastasis. We include both anti- and pro-tumor immune effects, in addition to the experimentally observed phenomenon of tumor-induced immune cell phenotypic plasticity. Using geometric singular perturbation analysis, we derive a rather simple model that captures the main processes and, at the same time, can be fully analyzed. Literature-derived parameter estimates are obtained, and model robustness is demonstrated through a sensitivity analysis. We determine conditions under which the parameterized model can successfully explain both metastatic dormancy and blow-up. Numerical simulations suggest a novel measure to predict the occurrence of future metastatic blow-up, in addition to new potential avenues for treatment of clinically undetectable micrometastases.

## 1 Introduction

Metastasis has long been viewed as the inevitable final step in cancer progression — the growth of a primary tumor leads to local invasion into the surrounding tissue where the tumor eventually encounters (or recruits) blood vessels, which provide it with a method for distant dissemination, which occurs stochastically downstream from the primary tumor. From this point of view it is reasonable to assume that treatment of a primary tumor at a *sufficiently early* time may have a good chance of preventing metastatic spread. However, recent evidence across multiple cancers has suggested that metastases can be seeded during early stages of primary tumor development, well before the primary tumor is clinically detectable [21,23]. Early micrometastases then may enter an extended period of *metastatic dormancy*, which can be as long as decades [24]. As a consequence of this early seeding and subsequent metastatic dormancy, the act of simply removing the primary tumor may be insufficient to effectively address metastatic disease.

In fact, the *opposite* observation has been made by researchers for more than a hundred years [23]: the removal of a primary tumor can trigger growth of previously unknown metastases throughout the body. We will use the term *metastatic blow-up* to describe this phenomenon of accelerated metastatic growth upon intervention at the primary site, but the process has also been referred to as the “stimulating effects of primary tumor resection on metastases” and “postsurgery metastatic acceleration” [3]. Although several theories have been proposed to explain metastatic blow-up, they have been — for the most part — qualitative and speculative.

Here we use the the *immune-mediated theory of metastasis* [47,49] to explain the poorly understood phenomenon of metastatic blow-up. The key of the immune-mediated theory is the inclusion of pro-tumor immune effects and immune cell phenotypic plasticity. In fact, there is ample literature (see a detailed discussion in [47]) showing that immune cells can be “re-programmed” or “educated” by the tumor to play a pro-tumor role. In [47] we developed a mathematical model for tumor-immune dynamics at a primary and a metastatic site, and we used it to analyze dynamics of metastasis such as metastatic growth at sites of injuries and the role of immune therapies. Here we simplify the model from [47] and focus on the tumor-immune dynamics at the metastatic site. Through the use of techniques from geometric singular perturbation analysis [25,27] we simplify the model to allow for the discovery of meaningful analytic results. We then parameterize our model using estimates from the literature, perform a parameter sensitivity analysis, and use the calibrated model to simulate clinically relevant scenarios. We find that the immune-mediated theory of metastasis can successfully explain metastatic blow-up in the case of highly inflammatory tumors and our model predicts that pro-tumor immune effects play a key role in the phenomenon. Based on our results, clinical tests to distinguish the makeup of local immune cell populations done only three to four weeks post primary resection may be capable of predicting whether or not metastatic blow-up will occur — even if it is only after an extended period of dormancy.

### 1.1 Metastatic Dormancy and Blow-up

Although metastatic dormancy and blow-up have been observed for many decades, the mechanisms underlying these processes have yet to be fully elucidated. Over the years, several potential mechanisms have been proposed, which we discuss briefly (see also[2,3,8,23]).

#### Theory 1: Resource monopolization by the primary tumor

Tumor growth is resource and nutrient intensive. Multiple tumor sites throughout the body compete with each other for these resources. The primary tumor — by virtue of being the primary — has priority access to these resources resulting in limited resources for the metastases and, consequently, limited metastatic growth. Upon primary tumor removal, the largest consumer of resources is eliminated, resulting in an increased access to nutrients for the metastases, resulting in metastatic blow-up.

An early, conceptually simple theory (also referred to as *atrepsis* theory [50]), resource monopolization has been rejected as a potential mechanism for dormancy and blow-up by Benzekry and collaborators in [3] wherein the corresponding mathematical model could not accurately fit experimental results performed in tandem.

#### Theory 2: Immune surveillance of metastases

Primary tumor growth elicits a cytotoxic immune response. If the primary tumor is sufficiently large, it may successfully evade this immune response, whereas the growth of smaller metastases may be effectively inhibited. By removing the primary tumor, the corresponding immune response is also abrogated. Without systemic immune surveillance, the previously controlled metastases are now capable of rapid growth, resulting in metastatic blow-up.

A number of mathematical models have appealed to this theory to explain their model results. Eikenberry et al. [14] used this reasoning to explain the blow-up they observed using a partial differential equation (PDE) model of (locally) metastatic melanoma. Similarly, the Enderling group [43,51] appealed to this framework to explain the results of their ordinary differential equation (ODE) model of immune trafficking between distant organs investigating potential mechanisms for *abscopal* effects [13] (the observation of systemic anti-tumor effects arising from a localized treatment). This theory has, however, been downplayed by Benzekry and collaborators [2,3] owing to the fact that previous experimental evidence (see [22] and references therein) has demonstrated that the phenomenon of *concomitant tumor resistance*, i.e. the suppression of metastatic tumor growth by a primary tumor, can occur in immuno-deficient mice.

#### Theory 3 Local promotion and global inhibition of tumor growth

Tumors are cytokine and chemokine factories [11] producing both *promoters* and *inhibitors* of tumor growth, and angiogenesis (among other effects). In 1993, Prehn [45] proposed that primary-tumor-produced *promoters* of angiogenesis act locally — supporting primary tumor growth — whereas angiogenic *inhibitors* act globally and thus inhibit metastatic growth (for a more recent review that discusses specific potential factors, see [8]). Therefore, metastases are expected to remain in a dormant state in the presence of a primary tumor, and grow upon the removal of the primary tumor and its associated systemic angiogenesis inhibitors, thus accounting for metastatic blow-up.

In [3], the authors explicitly include a generalized systemic inhibitor of tumor growth in their ODE model of distant tumor-tumor interactions and find that this model can most successfully fit experimental data compared to models of either resource monopolization or of angiogenesis inhibition. Similar results were reported in [23] where the authors demonstrated that the likelihood maximizing scenario that accounts for metastatic blow-up was suppression of metastatic growth when the primary tumor is present, and a significant increase to the metastatic growth rate upon primary tumor removal. Although compelling, the results from both of these works rely on highly generalized “inhibitors” and serve more as support for previously proposed theories and provide little in the way of concrete, testable predictions.

#### Theory 4: Primary resection induced inflammation

Relevant most specifically to blow-up, this theory invokes the well-known pro-tumor effects of the immune system and inflammation (see [1,37,47,49], for example) to explain the phenomenon. Primary tumor resection induces a significant systemic, but transient, inflammatory response [35,42], which produces pro-growth, proangiogenesis, and immune-suppressive factors associated with wound-healing. These pro-tumor factors enter the circulation and promote metastatic growth upon arrival to the sites of previously dormant micrometastases, yielding metastatic blow-up.

Previous models of metastatic dormancy and blow-up have neglected potential effects of inflammation. However, with accumulating evidence implicating the immune system in the metastatic process, the importance of including such effects is starting to be appreciated [2,47]. Primary resection induced inflammation is highly relevant in the clinical setting [46], and has yet to be thoughtfully explored using mathematical modeling. We address this shortcoming by developing a model of metastasic blow-up in the framework of the *immune-mediated theory* of metastasis [9,47,49].

### 1.2 Immune-Mediated Theory of Metastasis

The immune-mediated theory of metastasis posits that the immune system plays a key *pro*-tumor role in the metastatic process. A wealth of experimental and clinical evidence for this theory includes studies on

– metastasis to sites of injury [31,53]
– increased metastasis following primary resection [46] that can be inhibited, in some cases, with the use of non-steroidal anti-inflammatory drugs (NSAIDs) [17,28,36,46],
– increased metastasis upon biopsy [26],
– increased number and size of lung metastases in mice with a latent cytomegalovirus infection [56],
– immune preparation of the pre-metastatic niche (PMN) [29],
elimination of metastases with novel anti-inflammatory interventions [41],

and many more. Not only have *pro*-tumor immune effects been observed at each step of the metastatic cascade [37,47,49], but tumor-mediated immune cell phenotypic plasticity has also been documented [10,15,33,39,49], significantly complicating potential immune-based therapeutic interventions [47] and suggesting that metastasis may be more organized than previously thought. For further details concerning the *immune mediated theory* of metastasis, or immune involvement in metastasis more generally, please consult one of [4,9,37,47,49].

The theory of metastatic blow-up as a function of primary resection-induced inflammation (potential Theory 4 from the previous section) fits nicely within the framework of the *immune mediated theory* of metastasis. It is our goal to investigate the plausibility of this theory through the development, analysis, and simulation of a mathematical model for tumor-immune dynamics at a metastatic site.

### 1.3 Outline

The remainder of this work is organized as follows. Our model for tumor-immune dynamics at a metastatic site is developed in Section 2. Analysis of the model, including model reduction using geometric singular perturbation analysis, is done in Section 3. In Section 4 we parameterize the model, perform a sensitivity analysis, and investigate the model predictions with the help of numerical experiments. Simulation results are presented in Section 5, where we discuss a series of hypotheses about the relevance of the immune response and immune cell phenotypic plasticity on metastatic blow-up. Increased inflammation induced by surgical intervention as well as increased education dynamics have a strong effect on metastatic blow up. As a result we find that the prevention of significant resection-induced systemic inflammation and the reduction of immune cell phenotypic plasticity have beneficial effects on cancer control. In this context we briefly discuss the possibility of inducing metastatic blowup through primary tumor biopsies. Biopsies create small wounds, which can stimulate the immune response. While in most of our simulations a biopsy has a negligible effect, we found a small region in parameter space, where a biopsy can, in theory, lead to increase metastatic growth. Finally, a brief discussion of our results is presented in Section 6.

## 2 The Model

While several mathematical models have previously described metastatic dormancy and/or blow-up [3,14,23,51], they simplify, as all models do, the full extent of the problem. For example, Benzekry et al. [3] ignored immune effects and assumed that tumor cell loss was significantly higher at a secondary site *a priori* in order to accurately fit the experimental data. Although the extremely general model of Hanin et al. [23] demonstrates that metastatic blow-up is a robust biological phenomenon, it is that same generality that precludes any specific conclusions about the underlying mechanisms. In the models of both Eikenberry et al. [14] and Walker et al. [51], metastatic tumors had to be added to the simulations *manually* and post-resection inflammation was neglected. In order to relax these simplifications, we introduce a reduced version of the model for the immune-mediated theory of metastasis originally proposed in [47] with the goal of determining specific conditions necessary for metastatic blow-up.

### 2.1 The Original Immune-Mediated Model of Metastasis

The model that is the focus of this work is a reduction of a previously proposed model for the immune-mediated theory of metastasis [47]. In [47] we derived a eight-component ODE model for cancer growth dynamics at two sites, a primary tumor site and a metastatic site. At each site we consider the densities of tumorigenic tumor cells *u*, necrotic tumor cells *v*, cytotoxic immune cells *x* (CT-immune), and tumor educated immune cells *y* (TE-immune). The two sites communicate with each other through a free exchange of immune cells of both types, plus a seeding of tumorigenic cells from the primary tumor to the metastatic site. The eight-component model is quite complex, and a full mathematical analysis is not feasible. In [47] we used the model to show that tumor education of immune cells can explain tumor dormancy and blow-up, increased occurrence of metastasis at sites of injuries, and the unexpectedly poor performance of immune therapies [16].

Here we continue this line of research by focusing on the metastatic site. We simplify the model such that a full theoretical analysis is possible. This allows deeper insight into the dynamics of the immune-mediated theory. Using geometric singular perturbation theory we are able to find the root cause for metastatic dormancy and blow-up after primary removal. The key lies in the understanding of a one-dimensional dynamical system that lives on a slow-manifold of the system. In effect we reduce the original eight-component model [47] to a one-dimensional problem. Despite the significant reduction of complexity, the basic model features remain the same and the model can explain metastatic dormancy and blow-up.

In addition, we perform a sensitivity analysis for the three-component model of this paper to understand the relative importance of the various model parameters.

### 2.2 The Reduced Immune-Mediated Metastasis Model

Compared to the original model from [47] we make the following simplifying assumptions.

1. Rather than modeling the primary tumor site, we assume that a primary tumor exists and acts as a source of circulating tumor cell and TE immune cells.
2. We will assume that the only *explicit* pro-tumor immune effects are to increase the tumor cell growth rate. In their original model, Rhodes and Hillen included a second explicit role for TE immune cells — the inhibition of CT immune cell death. While no longer explicitly included in our model, CT immune cell impairment remains a part of the model via the tumor-mediated immune cell phenotypic plasticity.
3. Instead of modeling the necrotic cell compartment explicitly, we include this effect implicitely in the recruitment rate of the CT immune cells.

The end result of these simplifications is a basic model for the immune-mediated theory of metastasis,

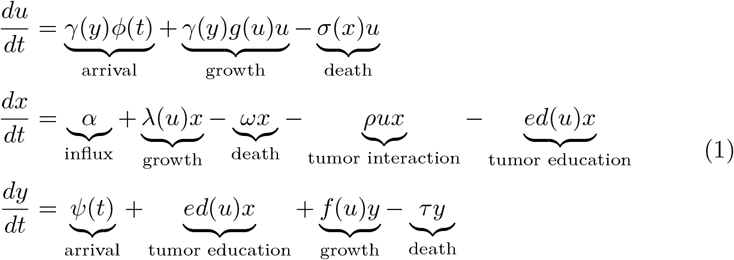

where *u*(*t*), *x*(*t*), and *y*(*t*) denote the tumor, CT immune, and TE immune populations at the metastatic site at time *t*, respectively. A schematic of this dynamical system is given in Figure 1. The underlying model assumptions are as follows:

– Tumor cells arrive at the metastatic site from the distant primary site at time-dependent rate *ϕ*(*t*), grow at rate *g*(*u*), and perish as a result of interactions with CT immune cells at rate *σ*(*x*). The term *γ*(*y*) represents the enhancement of both establishment and growth facilitated by the TE immune cells. Without sufficient evidence to the contrary, we have assumed that this TE immune cell enhancement is the same for both establishment and growth.
– CT immune cells arrive at the metastatic site with a constant influx rate *α*, and can be recruited by the tumor according to the function *λ*(*u*). Loss of CT immune cells can occur by one of (i) natural death, at rate *ω*, (ii) as a result of interaction with the tumor, at rate *ρ*, or (iii) as a result of tumor-education, at rate *ed*(*u*). The functional forms of *λ*(*u*), *ed*(*u*) will be discussed later.
– TE immune cells accumulate at the metastatic site in one of three ways: arrival from the primary site via the circulation at rate *ψ*(*t*), (ii) tumor education of a CT immune cell at rate *ed*(*u*), or (iii) local tumor-mediated recruitment according to the rate *f* (*u*). We also assume that TE immune cells perish at rate *τ*.

**Fig. 1.**
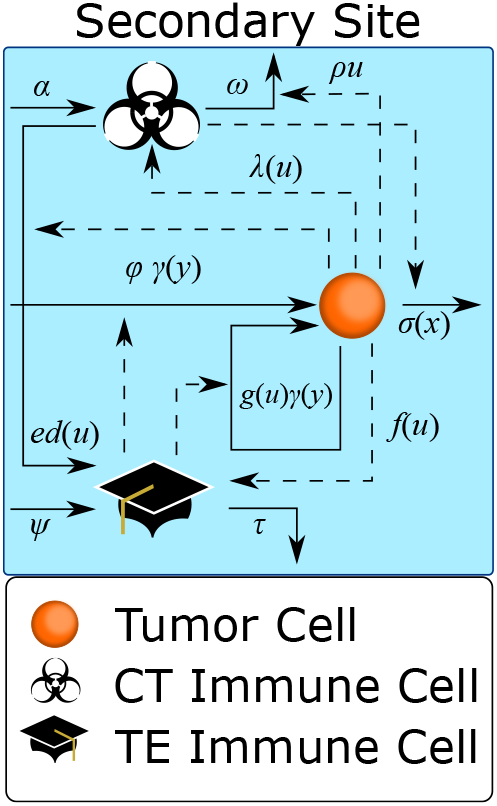
Cartoon model of the 3 ODE model of metastasis (1). Arrows indicate *positive* effects, and flat ends indicate *inhibitory* effects. Solid lines represent *direct* effects and dashed lines denote *indirect* influence. See text for details. Based on figure found in [47].

All parameters are assumed to be positive, and the functional coefficients are assumed to have the following behavior:

**Assumptions (A1):**

– The TE-immune enhancement factor of tumor growth and establishment, *γ*(*y*), is increasing from *γ*(0) = 1 to a finite, maximum value as *y* → ∞.
– the per-capita growth rate of the tumor cells *g*(*u*) is a decreasing, non-negative function with a carrying capacity *K* > 0, such that ∀*u* ≥ *K* we have *g*(*u*) = 0.
– The tumor cell death rate *σ*(*x*) is an increasing, strictly positive function.
– The immune recruitment rate *λ*(*u*) is an increasing and bounded function with *λ*(0) = 0
– The per capita growth rate of the TE immune cells *f* (*u*) is an increasing, bounded function such that *f* (0) = 0.
– The education rate *ed*(*u*) is an increasing function with *ed*(0) = 0.
– The immune influx rates *ϕ*(*t*) and *ψ*(*t*) are non-negative and bounded functions.
– Finally, all functions *γ*(*y*), *g*(*u*), *σ*(*x*), *f* (*u*), *ed*(*u*) are globally Lipschitz continuous in their argument.

For a full biological motivation of the above choices, we refer to the detailed discussion in our previous work [47]. Specific examples of these functional forms are as follows:

### 2.3 Examples of Functional Forms

Here we list some specific examples of the functional forms mentioned above, which will be used later in our numerical simulations.

– We assume that the metastatic tumor grows logistically [5,32,51] with intrinsic growth rate *r* and carrying capacity *K*,

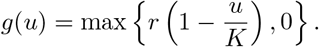
– TE immune cell enhancement of tumor growth *γ*(*y*) is modeled using an increasing hyperbolic tangent function [40,47],

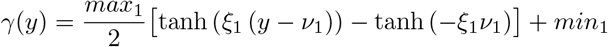

where 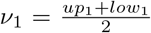 and 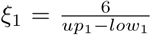 This function is a sigmoid curve that increases from 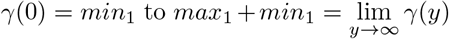. The parameters *low*_1_ and *up*_1_ are activation and saturation thresholds, respectively.
– Similarly, we use the same saturating functional form for the tumor cell death rate,

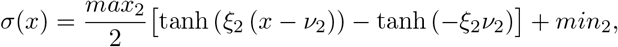

where 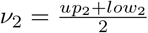 and 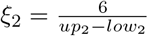.
– Tumor mediated expansion of the CT and TE immune cell populations are assumed to follow Michaelis–Menten kinetics [5,32,51],

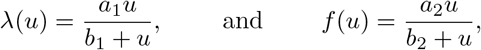

respectively, where *a*_*i*_ is the maximal recruitment rate and *b*_*i*_ is a half saturation constant, *i* = 1, 2.
– Tumor education of CT immune cells is assumed to be governed by the law of mass action [5],

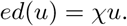

## 3 Analysis

This section is concerned with the analysis of the model of tumor-immune interactions at a metastatic site (1). The stability analysis of the three-component ODE model (1) is standard and we list the relevant results while skipping the detailed calculations. We then provide more details for the geometric singular perturbation analysis approach [25,27] in Section 3.3. Timescale arguments are used to reduce the model to a *slow manifold*. On the slow manifold we can then perform a more complete stability and bifurcation analysis (Section 3.4).

### 3.1 Basic Properties of the Immune-Mediated Model (1)

We summarize some elementary facts in the first Lemma.

#### Lemma 1

*Assume (A1). The model (1) has unique and bounded solutions that stay non-negative for finite time t* < ∞ *and for non-negative initial data.*

*Sketch of Proof.* Unique solutions exist globally because of the global Lipschitz continuity of the non-linearities. For non-negativity we show that along each of the three coordinate axes, the vector field, prescribed by our model equations, (1) is pointing into the positive octant.

To obtain uniform boundedness of solutions, we need stronger assumptions. We set

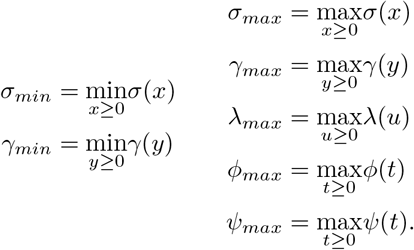

And we assume

**Assumption (A2):** *λ*_*max*_ < *ω* and *f*_*max*_ < *τ*.

#### Lemma 2

*Assume (A1) and (A2) then the solutions of (1) are uniformly bounded.*

*Proof* We first consider the *u* equation. Suppose that *u* ≥ *K*. Then, from (A1), *g*(*u*) = 0, and so the equation governing the evolution of *u* reads

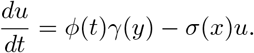

It follows that *u* will be decreasing whenever

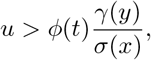

which is the case for

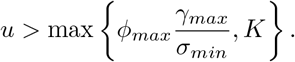

Hence *u* is bounded as

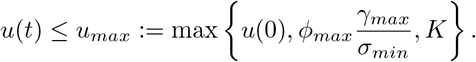

Next, we consider the *x* equation. Using the bound on *u* we find

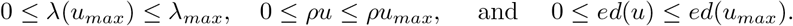

This leads to a relation between two new parameters *A* and *B*:

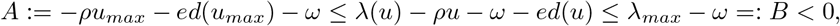

where the final inequality is by the assumption of the lemma. Therefore, we have an upper and lower bound on the ODE governing the *x* dynamics,

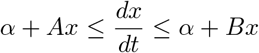

where *B* < 0. Hence *x*(*t*) is uniformly bounded.

Using the boundedness of both *u* and *x* allows us to reduce the *y* equation to a similar ODE as in the *x* case, yielding

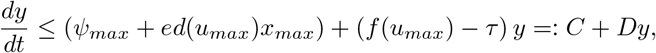

where *C* > 0 and *D* < 0 by the assumption of the lemma. Hence also *y* is uniformly bounded.

Assumption (A2) assures that the dynamics are bounded. It is a quite natural assumption, since it says that the maximal recruitment rates for the CT immune cells and for the TE immune cells are less than the natural death rates for these populations. Without any presence of cancer cells (as a source or at the metastatic site), CT immune cells reach a non-zero “healthy surveillance” steady state and the TE immune cell population drops to zero, corresponding to fully healthy tissue. Hence both immune cell compartments need stimulation from the cancer cells to expand their populations. We also note that these assumptions guarantee that both *λ*(*u*) − *ρu* – *ω* – *ed*(*u*) < 0 and *f* (*u*) − *τ* < 0 for all values of *u*. We now proceed to determine the steady states of the model (1).

### 3.2 Steady States

For simplicity, assume that the source terms, *ϕ*(*t*) and *ψ*(*t*), are constants, *ϕ* and *ψ*, respectively. While it may seem a strong assumption to have a constant source of both circulating tumor cells and TE immune cells, it is supported by the literature. First, in models of metastasis, shedding from the primary tumor is often modeled as a Poisson process [20,23] with strength proportional to primary tumor size. By assuming a threshold for the number of cells shed, a sufficiently large primary tumor will shed a relatively constant source of cells into the vasculature. Second, experimental models of metastasis have shown that (i) as many as 10^4^ cells per day can be shed from a primary tumor consisting of 10^8^ cells [12,54] and (ii) the vast majority of cells shed from the primary tumor successfully extravasate at a secondary site [6,34]. Therefore, it follows that a constant shedding would result in a constant (but scaled) establishment of metastases. As such, we believe that our assumption of a constant source of tumor and TE immune cells arriving at the metastatic site is well justified by the current literature. In anticipation of studying the effects of primary tumor interventions on the metastatic tumor, we will investigate two cases relevant for treatment, namely when the source terms are positive and when they are zero (corresponding to successful removal of the primary tumor).

**Case:** *ϕ* ≠ 0 and *ψ* ≠ = 0

Let 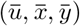 be a steady state equilibrium for the model (1). Based on the above assumptions (A1), (A2), we can formulate the steady state equations of (1) in the form of null-clines:

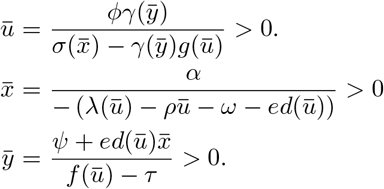

Steady states will be points, 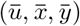, which lie on all three of the above nullclines. The number of such solutions is not immediately obvious, and depends on the choice of functional parameters. Further discussion of the number of steady states is done in a later section after specific choices have been made for the functional coefficients. Note that tumor extinction is impossible in this case — assuming that a source of tumor cells exists, the model predicts the persistence of metastatic disease.

**Case:** *ϕ* = 0 and *ψ* = 0

In this case we call the steady state 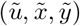. If 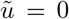 then we find an *extinction* steady state,

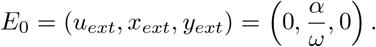

If 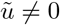 then our steady state 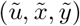 satisfies

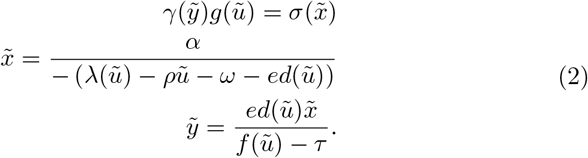

In general we have three defining equations for steady states, depending on the value of the source terms *ϕ* and *ψ*: a full-disease state, 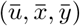, which exists when we have non-zero source terms, and without source terms there exists a disease-free state, (*u*_*ext*_, *x*_*ext*_, *y*_*ext*_), and a *persistent* disease state, 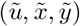. Discussion of the stability of the non-trivial steady states is delayed until the following section, where a timescale argument allows us to study stability much more easily. Before that, though, a simple computation yields the following stability result, which closely mirrors a similar result in the original two-site model [47].

#### Proposition 1

*Assume that ϕ* = 0 = *ψ and that* 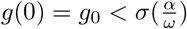. *Then the disease-free state, E*_0_, *is stable*.

*Proof* Evaluating the Jacobian of our system, *J*, at 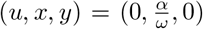 with *ϕ* = 0 = *ψ* gives us

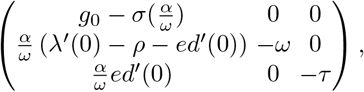

whose diagonal entries are its eigenvalues.

### 3.3 Timescale Reductions

In this section, we assume that the dynamics of the tumor are slow relative to those of the immune system. This assumption is biologically reasonable, as immune dynamics — such as immune response to an injury — occur on the timescale of minutes or hours [42], whereas tumor dynamics, especially dormant metastases, can be on the timescale of years or even decades [24]. Under the assumption of two timescales, we will perform quasi-steady state analysis of the model (1) using methods from geometric singular perturbation theory [25,27]. As we will demonstrate, this approach dramatically simplifies model analysis and allows us to gain significant insight into the underlying biology. We assume that the immune dynamics equilibrate quickly. In other words, we assume that all the parameters of the immune system equations in (1) are an order of magnitude larger than those of the tumor cell compartment. We introduce a small parameter *ϵ* > 0 and write

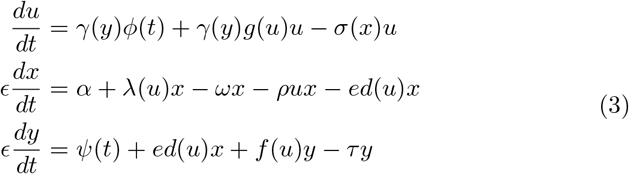

Using the notation *U* = *u*, *V* = (*x, y*) we write this system (3) in the more abstract form

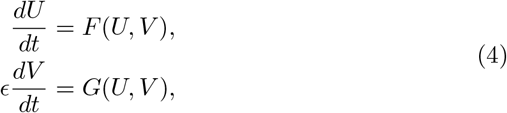

with

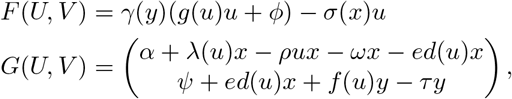

where this system (4) is called the *slow system*. We obtain the *fast system* by a transformation of the time variable as

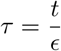

and obtain in the new variable

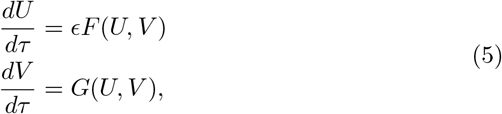

Letting *ϵ* → 0 the fast system (5) reduces to

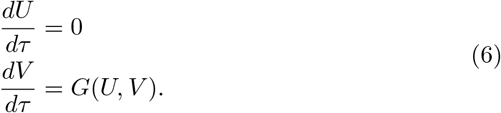

In other words, while the tumor density remains constant, the immune-components converge to their attractor. They cannot diverge to infinity, since the solutions are bounded (Lemma 2). The attractor of the immune dynamics is characterized by the set

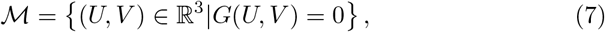

which is also known as the “slow manifold” of the system. The slow dynamics (4) for *ϵ* → 0 becomes

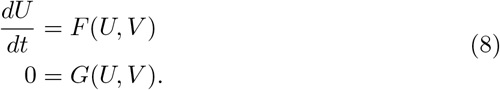

From the second equation in (8) we see that the dynamics “live” on the slow manifold 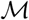, while the tumor dynamics on 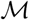 are given by the first equation of (8). Long term behavior of the system will be governed by the “slow” system (8). These observations are based on Fenichel’s geometric singular perturbation theory:

#### Theorem 1

(Fenichel [25]) *Suppose 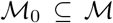 is compact, possibly with boundary, and normally hyperbolic, that is, the eigenvalues λ of the Jacobian* 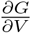 *all satisfy* 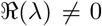. *Suppose F and G are smooth. Then for ϵ* > 0 *and sufficiently small, there exists a manifold* 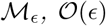 *close and diffeomorphic to* 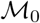 *that is locally invariant under the flow of the full problem (1).*

Figure 2 shows a plot of the null surfaces defined by setting 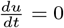 (purple), 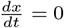 (blue), and 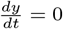 (red). The intersection of the immune-related null surfaces defines the slow manifold 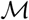 (green curve), while the intersections of all three surfaces (open circles) define the steady states of the system. We now provide a few results concerning the manifold 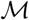.

**Fig. 2.**
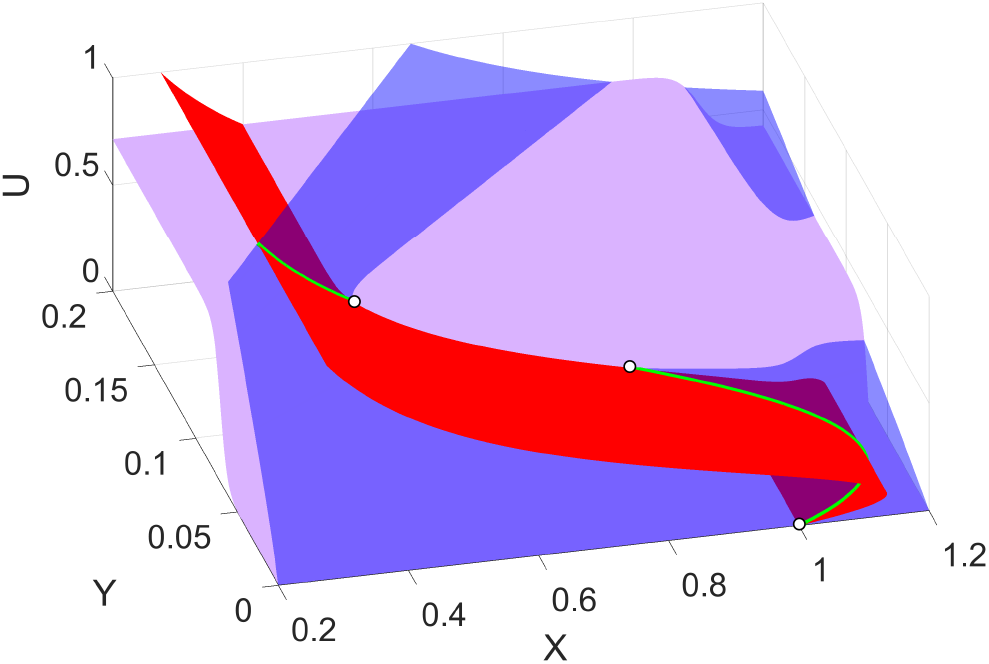
Null surfaces of the model (1). Blue is 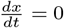, red is 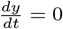 and purple is 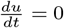. The slow manifold, 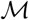, is the intersection between the *x* and *y* null surfaces, denoted by the green curve. Intersections of all three surfaces (denoted by circles) are steady states. Parameters as in Table 1, with the exception of *ϕ* = 0, and *ψ* = 0. Color figure available online.

#### Lemma 3

*The manifold* 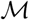 *is normally hyperbolic*.

*Proof* We compute the Jacobian

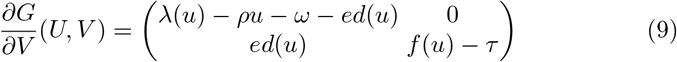

and note that the diagonal entries are both negative as a consequence of assumption (A2).

#### Proposition 2

*We can write* 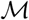 *as a graph*, (*u, x, y*) = (*u, x*(*u*), *y*(*u*)).

*Proof* Fix *u* ≥ 0. 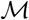 is defined via *G*(*U, V*) = 0. More explicity, this means that both of the following equations hold:

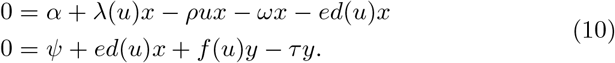

The assumption (A2) allow us to solve the first expression explicitly for *x*, yielding

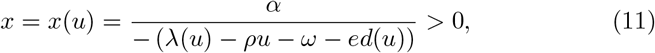

which is an explicit expression for *x* = *x*(*u*). Similarly for *y*, we can solve the second equation in (11) in terms of *u* and *x* as

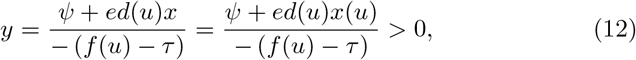

thereby giving us an explicit expression *y* = *y*(*u*).

Note that by Lemma 2, our dynamics will take place over a bounded domain in ℝ^3^, and so the functions *x* = *x*(*u*) and *y* = *y*(*u*) from Proposition 2 will be bounded and need only be considered over a bounded domain of R. Additionally, the continuity of *x*(*u*) and *y*(*u*) ensures that their images over a compact domain are themselves compact. Hence, 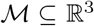 is compact. We are thus in the situation to apply Fenichel’s theorem to arrive at the following result.

#### Theorem 2

*For sufficiently small ϵ* > 0, *the dynamics of the reduced system (8) on the slow manifold* 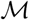 *provide a reasonable approximation (in the sense of Theorem 1) of the full system (4)*.

To understand the dynamics of the full system, we need only investigate the dynamics of the reduced system (8) on 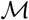, therefore, reducing the system of 3 ODEs in (1) to a single ODE in the tumor cell density, *u*,

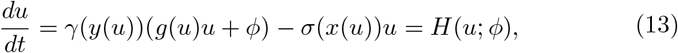

where *x*(*u*) and *y*(*u*) are defined as in equations (11) and (12), respectively. Consequently, the questions of number and stability of steady states reduce to the number of solutions to *H*(*u*; *ϕ*) = 0 and the sign of *H* on either side of these solutions, respectively. The stability of the steady states are simply determined by the sign of *H*(*u*; *ϕ*) on either side of the root, *u*_*_ : if *H*(*u*; *ϕ*) > 0 as 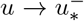, and *H*(*u*; *ϕ*) < 0 as 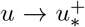, then *u*_*_ is stable, and it is unstable if the signs are reversed. Further study of the number and stability of steady states for our model is performed in the Section 3.4.

#### 3.3.1 Specific Example

As an example we use the functional coefficients introduced in Section 2.3 and arrive at the following description of the dynamics along the slow manifold 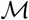 in terms of the derivatives 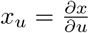 and 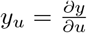, where *x*(*u*) and *y*(*u*) are as in (11) and (12), respectively.

##### Proposition 3

*Consider the functional forms from Section 2.3. Then on* 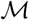 *we have that x_u_* > 0 *for u < u*_+_ *and x_u_* < 0 *for u > u*_+_, *where*

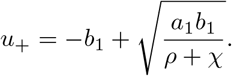

*Additionally, y*_*u*_ > 0 *for all* 0 ≤ *u* ≤ *K*.

*Proof* The proof follows from direct computation of the derivatives of *x*(*u*) and *y*(*u*) on 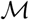. It is given in the Appendix for completeness.

Remark that the results of Proposition 3 — non-monotonicity of 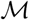 in the *x* coordinate and the monotonicity of 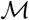 in the *y* coordinate — are highlighted in Figure 3. Initially, the *x* coordinate of 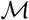 is *increasing* in *u*, but at the critical value of *u*_+_ ≈ 0.1, *x*(*u*) begins to *decrease*. On the other hand, it is clear in Figure 3 that *y*(*u*) is increasing throughout the domain of interest.

**Fig. 3.**
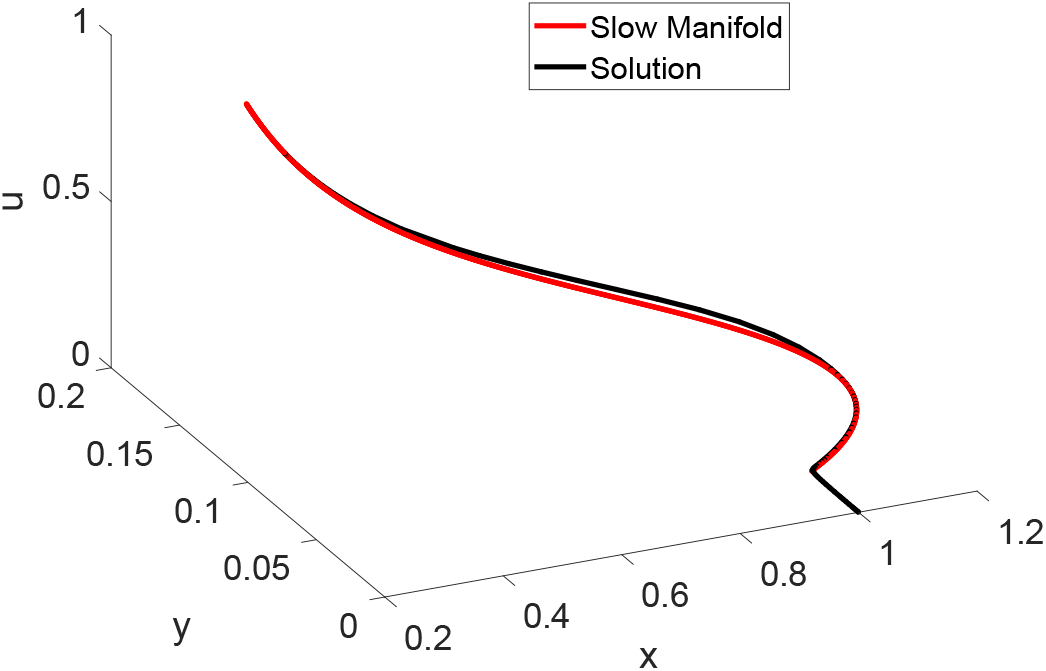
The slow manifold, 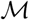 (red) and the solution to the system (1) (black). Initial conditions (*u*_0_, *x*_0_, *y*_0_) = (0, 1, 0). Parameters as in Table 1. Color figure available online.

### 3.4 Bifurcation Analysis

In this section, we exploit the approximation result of Theorem 2 to investigate bifurcations in two parameters: *ϕ*, the rate of circulating tumor cells (CTCs) arriving at the metastatic site, and *min*_2_, the minimum CT immune cell-mediated tumor cell death rate. These two parameters were chosen for use in our bifurcation analysis because of their relative sensitivities (see Figures 5 and 6) in addition to their clinical relevance. In particular, under the assumption that the primary tumor sheds tumor cells into the circulation at a rate proportional to its size [23], the parameter *ϕ* can be viewed as a measure for the size of the primary tumor. Furthermore, treatment of the primary tumor may decrease the value of *ϕ*. Combining these two observations allows us to interpret the effects of primary tumor growth and treatment on the metastatic tumor using bifurcation diagrams in the parameter *ϕ* (Figure 4, panel B).

**Fig. 4.**
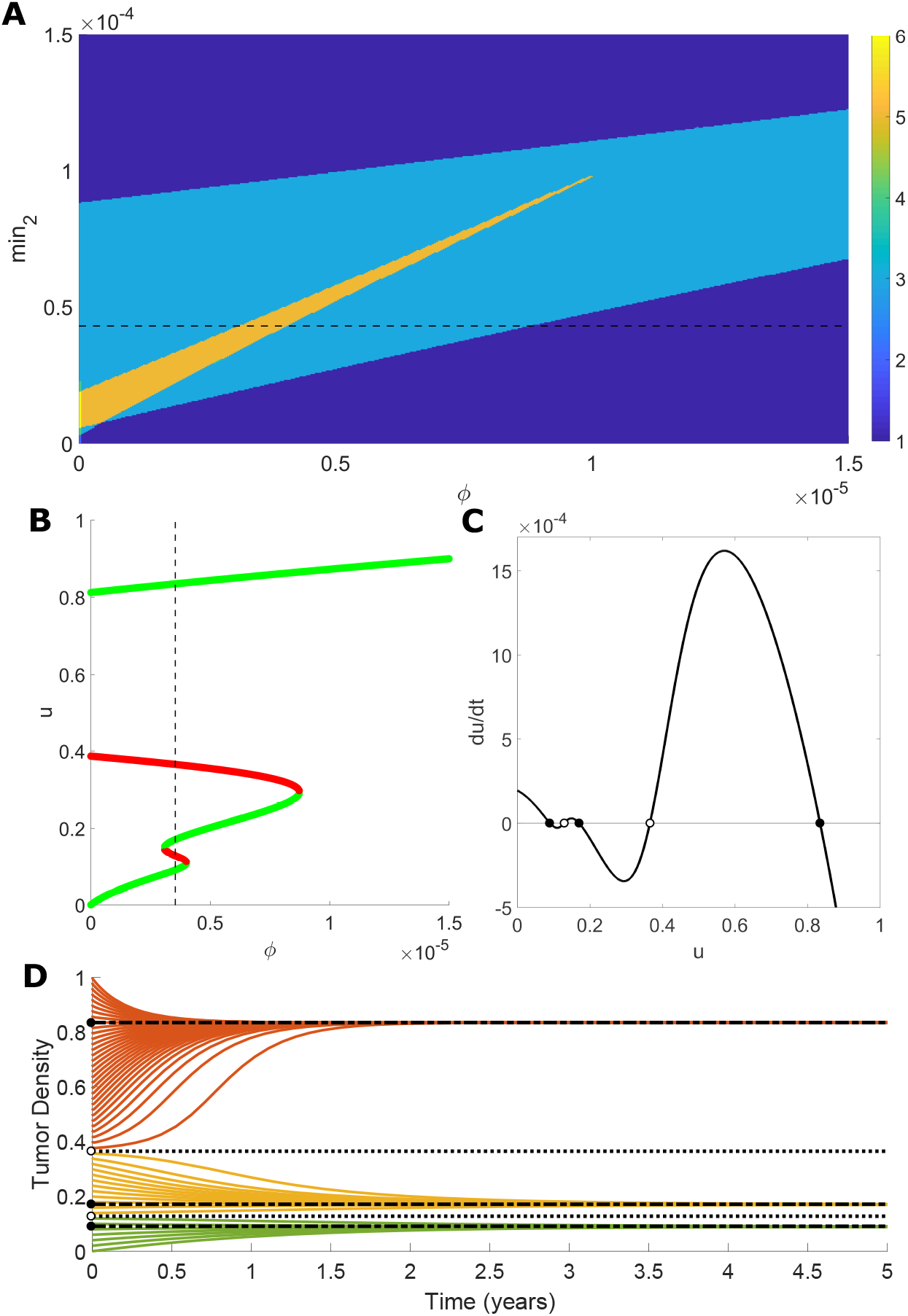
Confirmation of the validity of the slow-manifold approximations to describe the full system dynamics. (A) Two-dimensional bifurcation diagram in the parameters *min*_2_ (vertical axis) and *ϕ* (horizontal axis). Colors indicate the number of roots of *H*(*u*; *ϕ*) for different parameter pairs. Note that the maximum number of steady states is observed in the region along the leftmost vertical (corresponding to *ϕ* = 0) below approximately *min*_2_ ≈ 0.2. Horizontal dashed line denotes value of *min*_2_ = 4.255 × 10^−5^. (B) One-dimensional bifurcation diagram in *ϕ* for fixed value of *min*_2_ = 4.255 × 10^−5^. For each value of *ϕ* the corresponding steady state values are plotted. Stable steady states are colored green and unstable states are red. The vertical dashed line denotes value of *ϕ* = 3.5244 × 10^−6^. (C) Phase-line diagram using our slow-manifold approximation as described in Proposition 2 using fixed values of *min*_2_ = 4.255 × 10^−5^ and *ϕ* = 3.5244 × 10^−6^. Stable states are solid circles and unstable states are open circles. (D) (D) Solutions of the full model for 51 different initial tumor densities (initial conditions) lying in [0, 1]. Horizontal lines denote stable (dash-dot) and unstable (dot) steady state values. Trajectory color based on the longterm behavior. Parameters as in Table 1 with the exceptions of *max*_1_ = 0.2, *up*_1_ = 0.095, and *r* = 2.0 × 10^−4^. Color figure available online.

While a full characterization of all possible steady states is not available at present, we provide a brief discussion of the problem itself, and observed behavior of the model. Of interest is the number of solutions to equation (13)

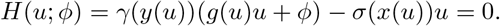

Observe that *H*(0; *ϕ*) = *γ*(*y*(0))*ϕ* > 0 and that the boundedness and positivity of *σ* and *γ* guarantee the existence of a sufficiently large value of *u* (call it *u*\*) such that *H*(*u*\*; *ϕ*) = *γ*(*y*(*u*\*))*ϕ − σ*(*x*(*u*\*))*u*\* < 0. Therefore, by continuity, we always have at least one positive steady state in [0, *u*\*]. Furthermore, the behaviors of *x*(*u*) and *y*(*u*) on the slow manifold 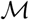 have been characterized (Proposition 3). Indeed, since *y*(*u*) is increasing in *u*, and *γ* is an increasing function, we know that the term *γ*(*y*(*u*))*ϕ* is increasing in *u*. In contrast, because *g*(*u*) is decreasing in *u*, the expression *γ*(*y*(*u*))*g*(*u*) may not be monotonic. We have determined that it begins at a positive value no smaller than *g*_0_ and evaluates to zero for *u* ≥ *K*. The final expression included in *H* is the death term, *σ*(*x*(*u*))*u*. While it is assumed that *σ* is decreasing, we know that *x_u_* changes from positive to negative at the value *u*_+_. The exact dynamics of the term *σ*(*x*(*u*))*u* therefore depend on the value of *u*_+_ and the choice of *σ*. The precise behavior is rather complicated and a full characterization is not feasible in general.

Despite this complexity, we can determine the number of solutions to *H*(*u*; *ϕ*) = 0 numerically for various choices of our bifurcation parameters. As can be seen in panel A of Figure 4, depending on our choices of values for the two bifurcation parameters, our model (with our specific choices of functional coefficients) has anywhere between one and six steady states (with the maximum six steady states observed in the bottom left corner of the presented plot). The boundaries between dark blue and light blue regions, as well as between light blue and orange regions represent saddle node bifurcations in which two steady states are created or destroyed. These bifurcations are illustrated in panel B, which is a bifurcation diagram in a single parameter (*ϕ*) for a fixed value of *min*_2_ (the dashed horizontal line in panel A). Returning to panel A, the bifurcations along the left vertical axis (near the horizontal axis) are associated with the stability of the disease-free state when the source terms *ϕ* = 0 = *ψ* (see Proposition 1). In contrast to our maximum value of six different steady states, the simple Kuznetsov model [32], upon which our model is based, was shown to have at most four different steady states. Therefore, our model modifications are responsible for the creation of two new equilibria.

In addition to the *number* of equilibria, the results from the previous section also allow us to determine the *stability* of these equilibria, as well. Panel C in Figure 4 is a phase-line diagram for our reduced system (13) on 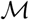, with fixed values of both bifurcation parameters (dashed horizontal and vertical lines in panels A and B, respectively), where solid dots denote *stable* steady states, and open dots represent *un*stable states. Confirmation of these approximation results are presented in panel D, where we present solutions to the full system (1) for different initial conditions, *u*(0) ∈ [0, 1]. It is clear in this plot that the stability determined using the reduced phase-line diagram in panel C is accurate, with trajectories moving toward stable states (dash-dotted) and away from unstable states (dotted). The accuracy of these approximated results — also illustrated in Figure 3 by the proximity of our solution to the computed slow manifold — justifies our analytical approach and will be of great benefit in Section 5.

## 4 Parameter Estimation and Sensitivity

Since several parameters are not available from the literature, we perform a sensitivity analysis, to be able to identify the most sensitive parameters. We find that some model parameters affect the early development of the solutions more than later effects, while other parameters become relevant only later. These results are demonstrated through a time-dependent sensitivity analysis, as shown in Figure 6. To our knowledge, this form of time-dependent sensitivity analysis has not been considered before, although it gives valuable information about the relative timing of the parameter sensitivities.

We begin by obtaining a baseline set of parameters informed by the currently available literature where possible (see Table 1).

**Table 1.** Model Parameters and the values used in presented simulations.

| Parameter | Description | Value | Units | References |
| --- | --- | --- | --- | --- |
| $r$ | tumor growth rate | $1.82 \times 10^{-4}$ | 1/time | [32,41] |
| $K$ | tumor carrying capacity | 1 | density | — |
| $min_1$ | min TE growth | 1 | — | — |
| $max_1$ | max (increase) TE growth | 2.5 | — | — |
| $low_1$ | growth activation | $5 \times 10^{-2}$ | density | — |
| $up_1$ | growth saturation | 0.1 | density | — |
| $\phi$ | CTC arrival | $1.85 \times 10^{-5}$ | density/time | [54] |
| $min_2$ | min death | $1.82 \times 10^{-4}$ | 1/time | — |
| $max_2$ | max (increase) death | $1.82 \times 10^{-4}$ | 1/time | — |
| $low_2$ | death activation | 1 | density | — |
| $up_2$ | death saturation | 1.2 | density | — |
| $\alpha$ | CT immune influx rate | $7.5 \times 10^{-3}$ | density/time | [32] |
| $a_1$ | CT expansion rate | $2.26 \times 10^{-2}$ | 1/time | [32] |
| $b_1$ | CT expansion damping | 0.404 | density | [32] |
| $\rho$ | fatal immune-tumor interaction rate | $3.11 \times 10^{-2}$ | 1/time | [32] |
| $\omega$ | CT decay rate | $7.5 \times 10^{-3}$ | 1/time | [32] |
| $\chi$ | immune education rate | $4.9 \times 10^{-3}$ | 1/time | [5,30] |
| $\psi$ | TE arrival | $8 \times 10^{-4}$ | density/time | [7,54] |
| $a_2$ | TE expansion rate | $1 \times 10^{-2}$ | 1/time | [32] |
| $b_2$ | TE expansion damping | 0.485 | density | [32] |
| $\tau$ | TE decay rate | $2.27 \times 10^{-2}$ | 1/time | [32] |
| $u(0)$ | Initial metastatic tumor density | 0 | density | — |
| $x(0)$ | Initial CT immune cell density | 1 | density | — |
| $y(0)$ | Initial TE immune cell density | 0 | density | — |

### 4.1 Parameter Estimation

Our choices of functional coefficients closely mirror those of our previous two-site model of the immune-mediated theory of metastasis [47]. Parameter estimates were obtained from the literature when available, and informed choices were made when no such estimation was possible. For the baseline parameters presented in Table 1, the following assumptions have been made. We have chosen the carrying capacity as *K* = 1 for simplicity. The growth rate *r* was estimated by fitting a logistic growth curve to normalized tumor growth data from [41]. The parameters *a*_1_, *b*_1_, and *ω* — shared by both our model and the Kuzntesov model [32] — were estimated by using the dimensional parameters presented in [32] and non-dimensionalizing as in [32] by assuming that the baseline CT immune and tumor cell densities are 10^8^ and 5 × 10^7^, respectively. These choices of baseline population values reflect a highly inflammatory metastatic setting with relatively few tumor cells compared to the primary site. The CT immune cell influx rate, *α*, was taken to satisfy *α* = *ω* in order to normalize the disease-free density of CT immune cells (this differs from Kuznetsov’s choice for *α*).

Threshold parameters *min_i_*, *low_i_*, *up_i_*, *max_i_ i* = 1, 2 have been estimated as follows. Assuming that a tumor population initiates a CT immune response, and that the immune system can effectively activate and destroy sufficiently small tumors, we have chosen 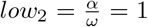, and *min*_2_ to be such that upon primary tumor removal, the disease-free steady state is stable (i.e. 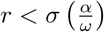: see also Figure 4). Although this assumption is made for our baseline parameter values, we eventually relax this assumption in the following sections when we are searching for conditions that will result in metastatic blow-up. The values of *up*_1,2_, *max*_1,2_, and *low*_1_ were chosen in order to have the associated rates change in time (i.e. the thresholds were passed at least once). We also remark that the choices made herein are in line with those used in [38].

Finally, the TE immune related parameters. The rate of tumor education of immune cells was informed by [5,30], but also chosen so that the tumor density *u*_+_, which corresponds to the value where *x*(*u*) is maximal on the slow manifold 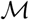 (see Proposition 3), was approximately 0.1. The growth and death parameters *a*_2_, *b*_2_, and *τ* were tuned from the Kuznetsov values to achieve a total immune population near the end of the simulation of approximately 1. The arrival rate, *ψ*, was estimated using data concerning shedding rates from the primary tumor [7], together with a likelihood of successfully arriving at the metastatic site [54]. Care was also taken in the parameterization procedure to assure that our solutions remained bounded (see assumptions A1 and A2). The results of this estimation process are summarized in Table 1.

### 4.2 Sensitivity Analysis

Using the parameters in Table 1 as our baseline parameters, we performed a basic parameter sensitivity analysis in order to determine the relative importance of the model parameters on the system outcome. A baseline solution for our tumor density, *u*, was obtained using the parameters in Table 1. For each model parameter, solutions were obtained for 30 different choices of the parameter value taken from the range ±10% of the baseline value. We report in Figure 5 the maximal change from our baseline solution in metastatic tumor density at time *t* = 840 days for each of our model parameters, with the maximum taken over all 30 tested values of each parameter. The endpoint time of *t* = 840 days was chosen so that our solutions had all reached steady state. Therefore, Figure 5 compares the effect of our parameters on the final steady state of the system.

**Fig. 5.**
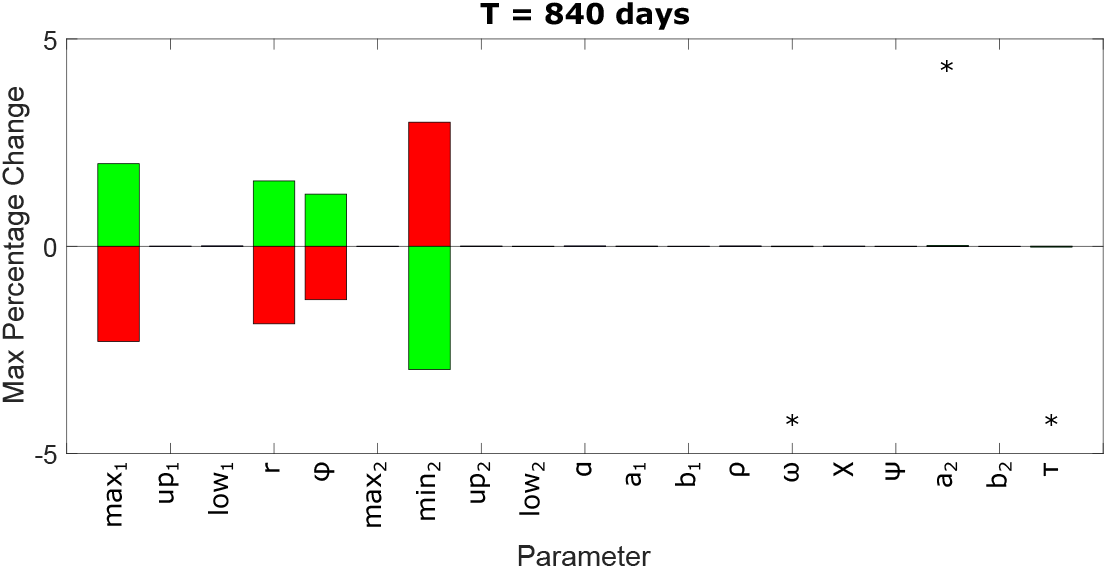
Maximum percentage change in tumor density compared to baseline at time *t* = 840 days. Red bars indicate that the resulting change in tumor density resulted from a *decreased* parameter value, while increased parameters are indicated by the green bars. Asterisks denote values that are of the order 10^−3^. Those without asterisks are smaller than 10^−4^. Further details in the text. Color figure available online.

Red bars indicate that the observed change occurred as a result of *decreasing* the parameter in question, while green bars indicate that an *increase* in that parameter was responsible for the observed change. As Figure 5 clearly demonstrates, there are only four model parameters that have any significant change on the model outcome: *min*_2_ (the minimum rate of CT immune cell mediated tumor cell death), *max*_1_ (maximum increase to tumor establishment and growth as mediated by TE immune cells), *r* (tumor cell growth rate), and *ϕ* (arrival rate of circulating tumor cells). All four of these parameters appear in the governing equation for tumor density. Furthermore, two of the four are threshold parameters; parameters for which we have scant experimental data to use in parameterization (see the previous section). The effect of a 10% change in all other parameters on the final tumor outcome is minimal — of the order 10^−3^ or less. Since *min*_2_ and *ϕ* are two of the most sensitive parameters in this setting, our choice to use them as bifurcation parameters in Section 3 (see Figure 4) is well justified. The results of Figure 5 provide an increased degree of confidence in our model predictions, as the end results are relatively robust to small perturbations in most of the model parameters.

### 4.3 Time-Dependent Sensitivities

Although the results of Figure 5 demonstrate the robustness of our final tumor density to small changes in parameter values, these particular results do little to elucidate the effects of perturbing the parameter values on the model *dynamics*. In order to address this particular shortcoming, we performed the same analysis (comparing tumor density of a perturbed solution to our baseline solution) once a week for 120 weeks, thereby giving us *time-dependent* sensitivities to perturbations in parameter values. The results of this extended sensitivity analysis are presented in Figure 6. Figure 6 shows the percentage increase (top) and decrease (bottom) for each of our model parameters (horizontal axis) over the course of 120 weeks (vertical axis). Percentages are indicated by the color bars. Before further analysis of these results, we will note that the abrupt horizontal line in both plots around the 35–40 week mark is a result of a significant change in tumor growth rate at about that time in the baseline solution (see panel (A) in Figure 7).

**Fig. 6.**
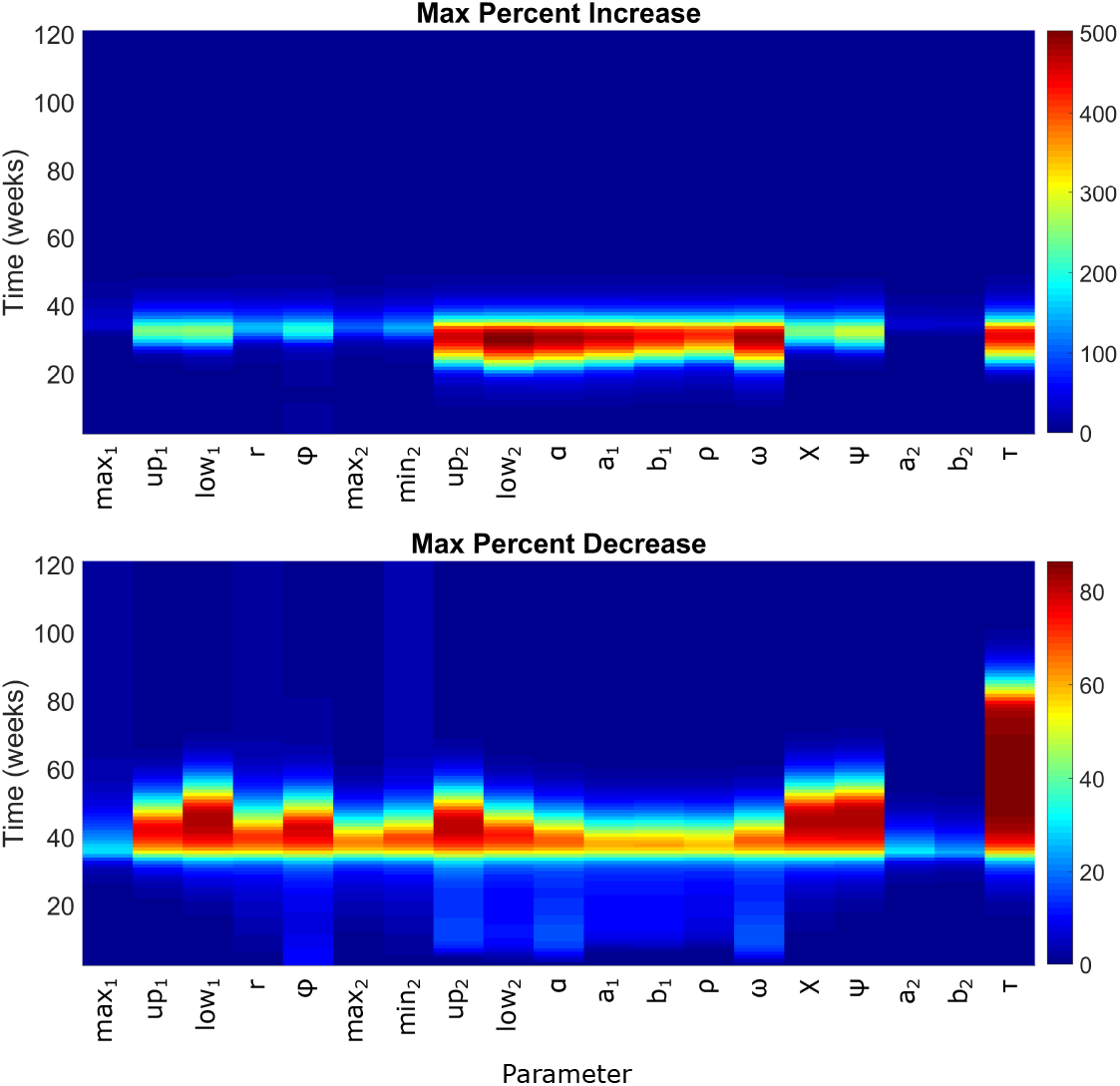
Maximum percentage change in tumor density compared to baseline at various times. Top is *increase* in tumor density compared to baseline, and bottom is *decrease*. Time is measured in units of weeks. Figure 5 represents the final time point in both top and bottom plots. Color figure available online.

**Fig. 7.**
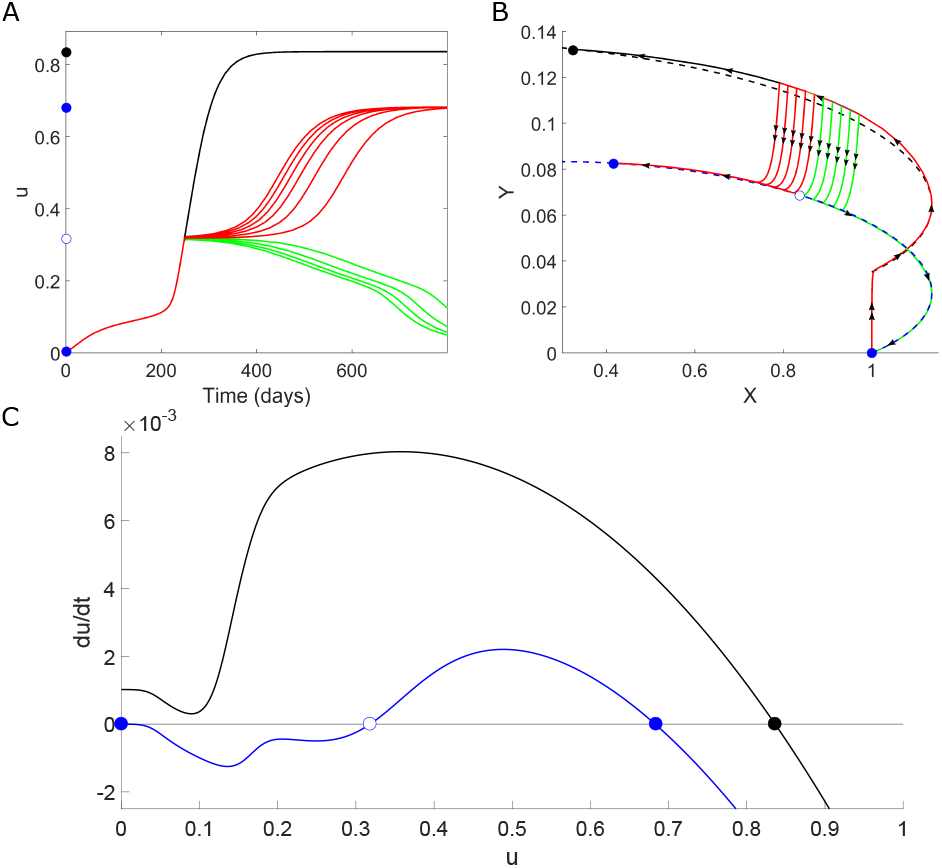
Results of simulated primary resection (P1). (A) The effect of primary resection on secondary tumor growth depending on resection time. Early interventions result in disease extinction (green) and late interventions result in disease persistence (red). (B) Model dynamics in 3-space, with the slow manifold for *ϕ*, *ψ* ≠ 0 is the dashed black curve, and the dashed blue curve denotes the slow manifold when *ϕ* = 0 = *ψ*. (C) Phase line diagrams for the cases when the source is on (black) and off (blue). Steady states are indicated by circles, solid representing stable and hollow representing unstable. The steady states are also marked in plots (A) and (B) for illustration. Parameters as in Table 1. Color figure available online.

Consider first the top plot. Most of the parameters have similar effects progressing through time: increasing until a point of maximal influence, followed by a period of decreasing influence. What this pattern tells us is *when*, relative to the baseline solution, the growth is most influenced by the parameter of interest. Take, for example, the parameter *low*_2_. For small times, we see little change from baseline. However, around the *t* = 20 weeks mark, we see the difference between baseline increase to a maximum of nearly 500% by *t* = 30. This period of increase reflects the fact that the baseline solution remains relatively unchanged over this period, whereas the perturbed solution is in a phase of rapid growth. The period of decreased effect compared to baseline (beginning at *t* = 30) is indicative of the perturbed system arriving at steady state, and the control system “catching up”. Finally, at long times when both solutions are in steady state, we see the effect of the parameter perturbation on the steady state (Figure 5 can be interpreted in this way). With this view, we can see that the effects of *low*_2_ are relatively early in disease progression compared to *ρ* or *ϕ*, and, despite its large peak, has little effect on the final outcome. Interestingly, the four most sensitive parameters by the end of the simulation all have relatively little effect during the transient phase of the model dynamics. Finally, we note that the effect of the establishment rate of tumor cells, *ϕ*, has two distinct peaks: an early one (almost immediately after the start of the simulation) and a later one that coincides with the peaks associated with most of the other model parameters.

Next, we consider the bottom plot in Figure 6. The dynamics here can be interpreted similarly, but this time with the results showing how much *smaller* the perturbed tumor is compared to baseline. Note that the length of the area colored yellow or red serves as a measure of the delay in tumor growth instigated by a change in the corresponding parameter. Thus, we see that the parameters *τ*, *ψ*, *χ*, and *low*_1_ are capable of the largest delays in tumor growth. Note that all of these parameters are associated with the pro-tumor TE immune cells. The four most sensitive parameters for long times cause relatively small delays in tumor growth. Additionally, the length of area colored yellow/red also serves as a measure of how “close” the perturbed system is to a bifurcation. Indeed, the dynamics in the TE immune cell clearance rate, *τ*, show an extended period of time wherein the perturbed system remains small compared to the baseline solution (large area of red extending from *t* = 35 weeks to *t* = 85 weeks), which indicates a close proximity of the phase line to the horizontal axis (see panel (C) in Figure 7) and therefore, a close proximity of the perturbed system to a bifurcation. For the perturbations considered here, no bifurcations were observed, but larger perturbations do see multiple bifurcations (see, for example, Figure 4). Finally, as noted in the previous case, *ϕ* has the earliest effect, and several parameters (*ϕ*, *up*_2_, *α*, and *ω*) now show the double peak dynamics discussed previously. Moreover, most of the parameters have non-trivial early effects, suggesting that there are multiple ways to inhibit early metastatic growth, with few of them having significant lasting effects.

## 5 Simulation of Clinically Relevant Cases

In this section we study some clinically relevant questions numerically. In Section 5.1 we show that the model can predict both metastatic growth and metastatic decline after resection of the primary. Section 5.2 scrutinizes several mechanisms to see which can explain metastatic blow-up. In Section 5.3 we argue that an observation of the immune population three to four weeks after surgery to remove the primary tumor can give valuable information about the progression of metastasis. Finally, in Section 5.4, we discuss treatment strategies to prevent metastatic blow-up based on our mathematical model.

### 5.1 Metastatic Growth or Decline

To begin a computational analysis of the above model and its dynamics on the slow manifold, we consider a case that shows both metastatic growth and metastatic decline after resection of the primary. The naive method of simulating primary resection is to set the source terms *ϕ* = 0 and *ψ* = 0, as done, for example, in [14,51]. This simulates the successful removal of 100% of the tumor and TE immune cells at the primary site and assumes no recurrence and no inflammation at the primary site.

The results for this simple case of primary resection are presented in Figure 7. Panel A shows the metastatic tumor density as a function of time in the cases of no (black curve), early (green curves), and late (red curves) primary resections. If intervention at the primary site is performed sufficiently early, the metastatic tumor goes extinct (green). If, however, the primary tumor is left untreated for too long, the metastatic tumor grows, resulting in metastatic persistence (red).

Panel B shows two slow manifolds as dashed lines, pre-resection in dashed black and post-resection in dashed blue. All solutions begin at (*x, y*) = (1, 0) and quickly move onto the pre-treatment slow manifold (dashed black). At resection, the orbits transition from the dashed black to the dashed blue manifold, which has two additional equilibria — an extinction state and an unstable saddle point (open circle). Depending on which side of the saddle the orbits land, they will either go extinct (green curves) or persist (red curves). Note that the transition along the saddle point can take quite some time, giving one possible explanation for *metastatic dormancy*. Panel B also shows the close alignment of the solutions (colored solid lines) with the slow manifolds (dashed lines), confirming our scaling approach made earlier. Panel C shows the corresponding phase-line diagrams for the control (black) and post-resection (blue) parameters on the corresponding slow manifolds, 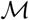.

Even though this process allows for metastatic growth post-resection, the resulting metastasis is smaller than the control case (red lines compared to black line in panel A). Hence metastatic blow-up is impossible in this case.

### 5.2 Metastatic Blow-Up

We are interested in determining conditions that result in metastatic blow-up upon primary intervention, which we define as follows:

#### Definition 1

We say that the model predicts *metastatic blow-up* if primary intervention eventually leads to a significantly larger metastatic tumor as compared to the case of no intervention at the primary site.

We saw already that the previous case, analyzed in Section 5.1 cannot lead to metastatic blow-up. In fact, if the pre-intervention system has only one stable equilibrium, the results from the previous section show that blow-up is impossible as the removal of the source of CTCs decreases the value of the largest possible stable state. As a consequence, to observe metastatic blow-up we require bi-stability in our pre-resection system, to allow for one small dormant state and a larger full disease state. For the following simulations we chose parameters that allow for this bi-stability, and which are in the biologically relevant range. More specifically, we changed two parameters from those presented in Table 1: *up*_1_ = 0.2 and *r* = 2.0 × 10^−4^. The black curves in Figure 8 show the dynamics of this newly parameterized model. The black phase-line on the right shows two stable attractors (solid black circles), separated by an unstable saddle point (open circle). Due to its importance in shaping the overall model dynamics, we will denote by 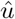 the value of this unstable saddle point, and refer to it as the *blow-up threshold*.

**Fig. 8.**
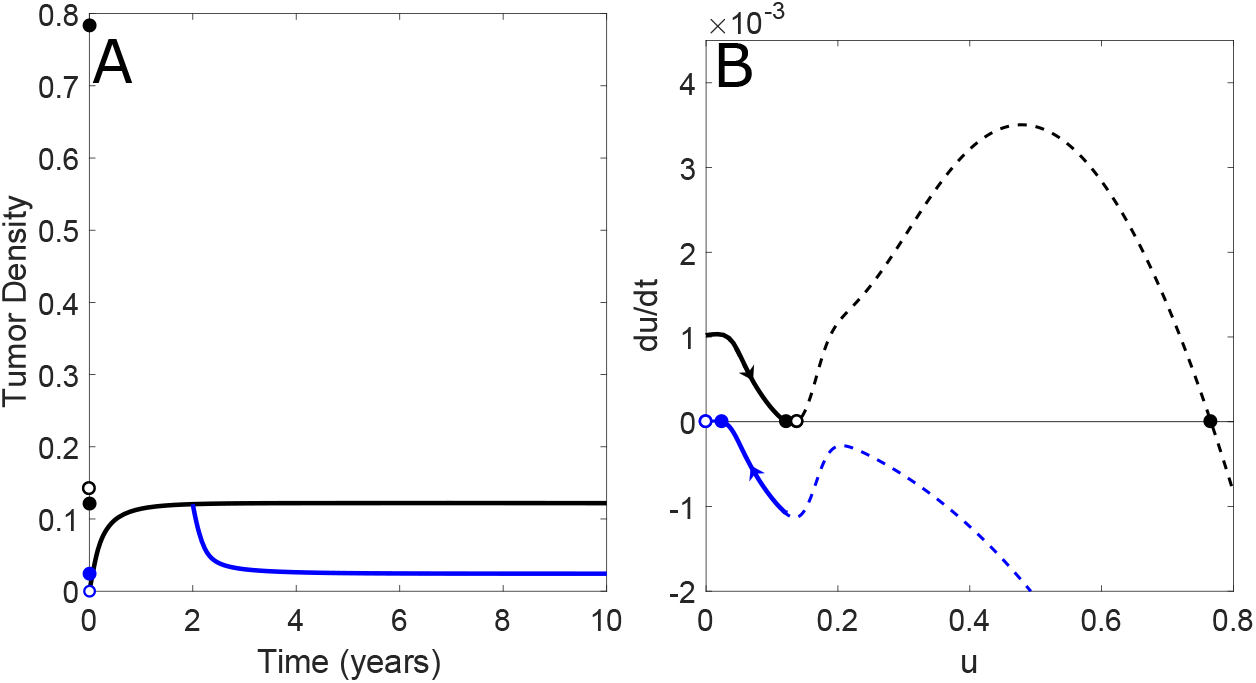
Case (M1): Simple simulation of primary resection. In both panels, primary resection is simulated by setting the source terms *ϕ* = 0 = *ψ* after two years. (A) Time dynamics of metastatic tumor density for a control case (black curve) and in the case of primary resection (blue curve). The dots on the vertical axis correspond to the steady states and their stability in the control (black) and post-resection (blue) cases. (B) Dashed lines are the phase-line diagrams for the control (black) and post-resection (blue) cases. Solutions to the full model are the solid lines superimposed, with direction of movement indicated by the arrows. Parameters as in Table 1, with the exceptions of *up*_1_ = 0.2 and *r* = 2.0 × 10^−4^. Color figure available online.

In order to analyze the ability of our model to reproduce metastatic blow-up, we consider individual mechanisms in increasing order of complexity:

**(M1) Naive Primary Resection:** Here we consider the naive primary resection model, where 100% of the primary tumor is removed, and no inflammation is induced. We use this case as a null-hypothesis. Primary resection is modeled by setting the two influx terms equal to zero: *ϕ* = 0, *ψ* = 0.
**(M2) Primary Resection and Systemic Inflammation:** Primary resection is an invasive surgical procedure which produces a significant, yet transient, systemic inflammatory response. For example, serum interleukin (IL) 6 concentrations have been shown to increase upwards of 500 times compared to basal levels in response to open curative resection for colonic cancer [42]. Even the less invasive primary tumor *biopsy* can be responsible for systemic inflammation (measured by presence of neutrophils in lung airways in a murine model of metasatic breast cancer) of strength 3–4 fold basal levels [26]. The application of NSAIDs can significantly reduce the risk of short-term metastatic blow-up in breast cancer [17,46]. In this second model of primary tumor resection, in addition to the naive simulation of primary resection (M1) we include a transient inflammatory response at the secondary site by multiplying the number of CT immune cells at the metastatic site at the time of intervention by a factor *θ* ≥ 1 (referred to as the “inflammation level”). This results in a short spike in CT immune cells that rapidly dissipates in the space of a few days to a week (depending upon the inflammation level simulated), matching timescales reported in the literature [17,26,42,46].
**(M3) Primary Resection, Systemic Inflammation, and Increased Tumor Pro-tumor Immune Presence:** Based on the results of Benzekry [3] and Hanin [23], suggesting that the growth rate of metastases *increases* upon primary resection, we investigate potential mechanisms within our model that will allow us to reproduce these results. The major role of pro-tumor TE immune cells in our model is through the growth-enhancement function, *γ*(*y*). We postulate that the population of TE immune cells *grows* in response to the primary tumor resection. Such a hypothesis, while novel, is in the same spirit of previous suggestions (summarized in Section 1). Within our modeling framework, there are three separate ways to increase the TE immune cell population after primary resection:

**(M3a)** *Increasing* the tumor-mediated recruitment rate, *a*_2_.
**(M3b)** *Decreasing* the death rate of TE immune cells, *τ*.
**(M3c)** *Increasing* the rate of tumor-education of CT immune cells, *χ*.
**(M4) Biopsy-induced Inflammation:** As a slight tangent, we also use our model to simulate the effects of a primary tumor biopsy on the growth of a metastatic tumor. The simulation is similar to that described in (M2), with the exception that we do not turn the source terms to zero, and, in order to closely match previously reported results, the inflammation level simulated is significantly lower than in the case of primary resection.

We want to investigate if any or all of the above processes can explain metastatic blow-up. A summary of the results is given in Table 2.

**Table 2.** Summary of the ability of different mechanisms to reproduce metastatic blow-up

| Mechanism | Figure | Dormancy | Blow-up |
| --- | --- | --- | --- |
| (M1) Naive Resection | 7, 8 | yes | no |
| (M2) Resection and Systemic Inflammation | 9 | no | no |
| (M3) Resection, Inflammation, and Increased TE immune cells | — | — | — |
| (M3a) Increased Recruitment | — | no | no |
| (M3b) Decreased Clearance | — | no | yes |
| (M3c) Increased Education | 10 | yes | yes |
| (M4) Biopsy-induced Inflammation | 11 | no | yes |

#### (M1) Naive Primary Resection

In this and all of the following simulations (unless noted otherwise), primary resection is performed two years from the beginning of the simulation. This date is chosen so that the metastatic tumor has reached a steady state. The blue lines in Figure 8 show the resection case. We see in the phase-line plot on the right, that upon resection, the metastasis declines to a very small positive value, hence in this case neither metastatic blow-up, nor metastatic growth is possible after resection of the primary.

#### (M2) Primary Resection and Inflammation

Figure 9 illustrates the results of including an inflammatory response (as previously described) to primary resection. Panel A shows the metastatic tumor density as a function of time in the case of no primary intervention (black) and in the case of primary resection (*ϕ* = 0 = *ψ*) with various inflammatory responses, ranging from *θ* = 2 times baseline to *θ* = 500 times [42] (green to red curves). Although there is a short period of inflammation-induced metastatic growth for sufficiently large inflammatory responses (panel C), in all cases the growth arrests and the tumor eventually shrinks to a small, positive steady state value.

**Fig. 9.**
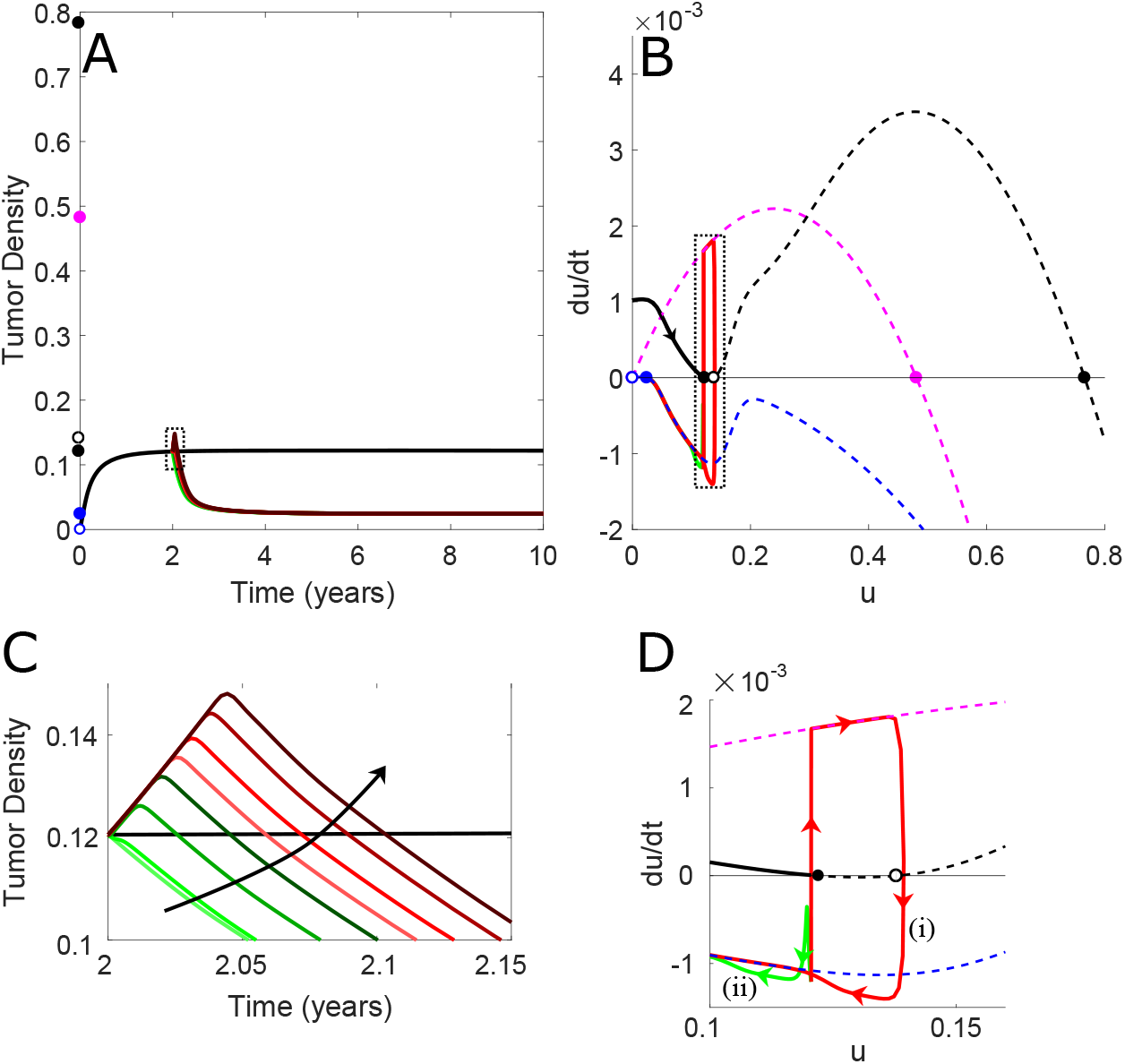
Case (M2): Simple resection with transient period of inflammation. In all panels, primary resection is simulated at the two-year mark by setting the source terms *ϕ* = 0 = *ψ* and by increasing the density of CT immune cells at the metastatic site by 2, 5, 10, 25, 50, 100, 250, or 500 times (arrow in panel C indicates the direction of *increasing* simulated inflammation). (A) Time dynamics of metastatic tumor density for a control case (black curve) and in the case of primary resection with various strengths of post-resection inflammation (green and red curves). The dots on the vertical axis correspond to the steady states and their stability in the cases of control (black), inflammatory (magenta), and post-resection (blue) parameters. The region enclosed in the dashed box is enlarged in panel (C), where the arrow indicates the direction of *increasing* inflammation. (B) Dashed lines are the phase-line diagrams for the control (black), inflamed (magenta), and post-resection (blue) parameters. Solutions to the full model are the solid lines superimposed, with direction of movement indicated by the arrows. The boxed region is depicted in panel (D), where red (i) and green curves correspond to inflammation levels of 100 and 2 times, respectively. Parameters as in Table 1, with the exceptions of *up*_1_ = 0.2 and *r* = 2.0 × 10^−4^. Color figure available online.

We can again exploit the slow manifold approximation of the full system to understand what is responsible for these dynamics (panels B and D). In panel B we show the phase-line diagrams for the control case (black dashed), for the resection case (blue-dashed) and for the highly inflamed case (dashed magenta). The highly inflamed line corresponds to the dynamics when the saturating functions *γ*(*y*) and *σ*(*x*) are saturated near their maximal values.

Panel D shows that for sufficient levels of resection-induced inflammation, the dynamics jump from the control state (black) to the highly inflamed state (magenta), during which time the metastatic tumor grows. Once the inflammation subsides, the solution drops to the post-resection state (blue) and decays to a small metastatic tumor. Even if the inflammation-induced growth is sufficient to push the metastatic tumor density into the basin of attraction of the larger control steady state (red curve), the fact that the primary tumor has been removed (and so *ϕ* = 0 = *ψ*) means that the dynamics governing metastatic growth *post*-resection are different than they were *pre*-resection, and the only stable equilibrium is a small, positive value.

Taken together, the results from Figure 9 demonstrate that removing the source terms has a stronger effect on the post-resection dynamics than a brief period of inflammation-mediated growth can overcome. This case does not support metastatic blow-up.

#### (M3) Resection, Inflammation, and Increased TE Immune Cells

It is in this step that we must leave the realm of experimentally or clinically documented effects, and begin exploring potential biological mechanisms proposed by our mathematical model. We consider the three cases (M3a), (M3b), and (M3c) each conditioned on the requirement that any saddle point that appears in the phase-line diagram *post*-resection must coincide with the original blow-up threshold, 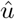. This requirement limits the number of admissible solutions, and allows for a meaningful comparison of the three treatment cases. For simplicity, we will refer to this condition as “the geometric condition.” Panel C in Figure 10 illustrates this condition in the case of increasing the parameter *χ*: the leftmost equilibrium is shared between the pre-resection (black) and the post-resection (blue) curves.

**Fig. 10.**
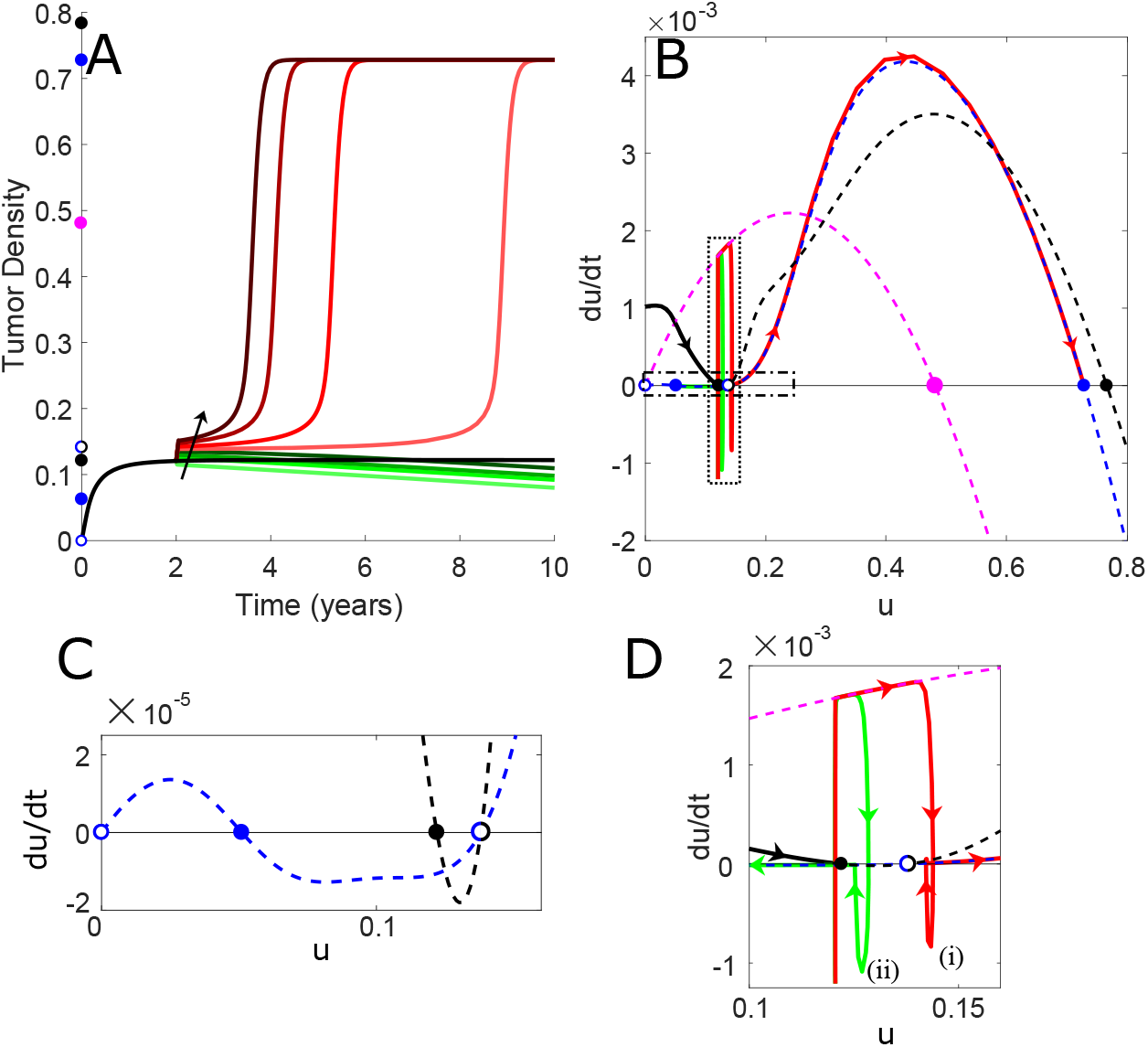
Case (M3c): Resection with transient inflammation and increased immune cell education rate post-resection. In all panels, primary resection is simulated at the two-year mark by setting the source terms *ϕ* = 0 = *ψ*, by increasing the density of CT immune cells at the metastatic site by the times indicated in Figure 9 (arrow in panel A indicates the direction of *increasing* simulated inflammation), and by increasing the rate of immune cell education, *χ*, by a factor of 2.42. (A) Time dynamics of metastatic tumor density for a control case (black curve) and in the case of primary resection with various strengths of post-resection inflammation (green and red curves). The dots on the vertical axis correspond to the steady states and their stability in the cases of control (black), inflammatory (magenta), and post-resection (blue) parameters. (B) Dashed lines are the approximate phase-line diagrams for the control (black), inflamed (magenta), and post-resection (blue) parameters. Solutions to the full model are the solid lines superimposed, with direction of movement indicated by the arrows. The region within the vertical box (dotted lines) is depicted in panel (D), where red (i) and green (ii) curves correspond to inflammation levels of 100 and 2 times, respectively. The horizontal box (dash-dotted lines) in panel (B) is depicted in panel (C), which depicts the pre- (black) and post-inflammation (blue) dynamics only. Parameters as in Table 1, with the exceptions of *up*_1_ = 0.2 and *r* = 2.0 × 10^−4^. Color figure available online.

**(M3a):** In order to satisfy the geometric condition in case (a), we increase the tumor-mediated recruitment rate of TE immune cells *a*_2_ by a factor of 7.35. Such a large increase results in the violation of the assumption (A2), resulting in unbounded solutions (results not shown) and leading us to reject this case as a potential mechanism for metastatic blow-up.

**(M3b):** Decreasing the TE immune cell death rate from *τ* to 0.3798 *τ* satisfies the geometric condition for case (b). Resulting model solutions remain bounded and metastatic blow-up is possible for sufficiently strong inflammatory responses to the primary resection. However, the *rate* of blow-up does not match clinical observations. Retsky et al. [46] report two peaks in breast cancer relapse — a pronounced *early* peak occurring approximately 18 months after treatment, and a broader *late* peak beginning approximately four years after treatment and extending to over 15 years post-treatment. In contrast, the blow-up observed as a result of mechanism (b) occurs within at most 6 months (results not shown) — much too soon post-resection to be responsible for the clinical observations.

**(M3c):** Finally, the geometric condition is satisfied via mechanism (c) by increasing the rate of tumor education of immune cells, *χ*, by a factor of 2.42 (panel C in Figure 10). In this case, not only do solutions remain bounded, but metastatic blow-up is possible for sufficiently large inflammatory responses and the *rate* of blow-up also much more closely matches the clinical observations of Retsky et al. [46] (panel A in Figure 10). Panel D shows that the saddle point (open circle) acts like a threshold, determining the final outcome of the metastatic tumor: solution trajectories that fall from the highly-inflamed magenta curve onto the post-resection blue curve to the *left* of the threshold decay to a small, dormant tumor (green curve), whereas those that fall to the *right* (red curve) will eventually result in metastatic blow-up. In this setting it is clear why the terminology *blow-up threshold* has been adopted for this unstable saddle point 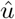. The time of dormancy is determined by a slow transition along this saddle point. It is because of the relative flatness of the post-resection curve that extended periods of dormancy before metastatic blow-up are possible.

#### (M4) Biopsy-Induced Inflammation

Although generally harmless, there is evidence to suggest that the inflammation induced from a biopsy of a primary tumor can lead to an increased incidence of metastasis [26]. Therefore, as a final application of our model in this section, we simulate a small jump in inflammatory cells at the metastatic site in response to a primary tumor biopsy (simulated as previously described), and investigate under what conditions this can result in metastatic blow-up. It is important to note that in these simulations, we have not simulated primary resection, and so we leave the positive source terms *ϕ* and *ψ* untouched.

The results of this investigation are summarized in Figure 11. Whereas the right plot is a bifurcation diagram in the parameter *ϕ*, the left plot shows the outcome at the metastatic site for different combinations of the parameter *ϕ* and the level of inflammation *θ* incurred from the biopsy at the primary site (vertical axis). The maximum inflammation level was chosen to be ten times basal levels to match the order of magnitude reported in the literature [26].

**Fig. 11.**
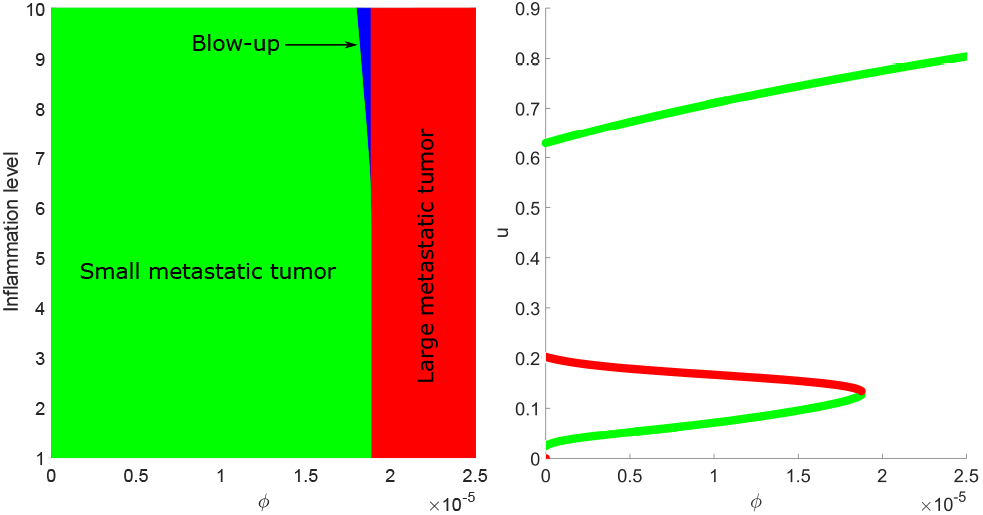
Case (M4): Effect of primary inflammation on metastatic tumor. (Left) Metastatic outcomes for pairs of inflammation level *θ* (vertical axis) and arrival rate of CTCs *ϕ* (horizontal axis). Portions of parameter space colored green indicate situations in which biopsyinduced inflammation has little to no effect on long term dynamics, and the metastatic tumor remains small. Regions in red indicate parameter values for which a large metastatic tumor is expected independent of any primary intervention. Parameters in the blue region result in metastatic blowup in response to the inflammation at the primary site. (Right) Bifurcation diagram in the source of CTCs, *ϕ*, illustrating the number and value of steady states for different values of *ϕ*. Green steady states are stable, whereas red states are unstable. Parameters as in Table 1 with the exceptions of *up*_1_ = 0.1 and *r* = 2.0 × 10^−3^. Color figure available online.

For small values of *ϕ* (corresponding to small, or early-stage primary tumors), the system exhibits bi-stability (right, green branches). However, as the initial metastatic tumor population is always assumed to be *u*(0) = 0, the system will always tend towards the smaller of the two stable steady states

– indicated in the left figure by the large green region. As *ϕ* increases past a bifurcation point at approximately *ϕ*\* ≈ 1.9 × 10^−5^, this bi-stability is lost, and only one steady state (corresponding to a large metastatic tumor) remains
– indicated in the left plot by the region colored red. For a small window near the bifurcation value of *ϕ*\*, however, a third outcome is possible: metastatic blow-up. If a sufficiently strong inflammation is triggered (*θ* large enough), then the metastatic tumor can grow *beyond* the unstable node (red branch on the right) and find itself in the larger steady state’s basin of attraction, resulting in a larger metastatic tumor compared to the case of no intervention at the primary site (i.e. metastatic blow-up). Hence systemic inflammation induced from even a small intervention at the primary site (such as a biopsy) is a potential mechanism to jump from a small metastasis to a large one. However, the region in parameter space where this can happen is rather small.

### 5.3 Immune Cells as Diagnostic Tool

In the previous section we identified a mechanism for metastatic blow-up: an increased metastatic growth triggered by the inflammation resulting from primary tumor resection and sustained by a post-resection increase of tumor-educated immune cells. Based on this process we investigate the consequences and potential benefits for diagnosis and treatment.

A critical parameter in case (M3c) was the blow-up threshold 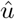, corresponding to the *u*-value of the saddle point on 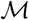. By Proposition 2, the manifold 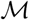 is a graph on *u*, hence we can express the blow-up threshold in terms of the immune population 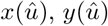 at the metastatic site. The advantage of this approach lies in the relative simplicity of measuring local immune cell populations at likely locations of metastatic disease compared to the difficulty of detecting micrometastases. Figure 12 shows the fraction of immune cells at the metastatic site that are CT for a control scenario (black curve) in addition to the primary resection scenarios investigated in for mechanism (M3c). The dashed horizontal line denotes the blow-up threshold of

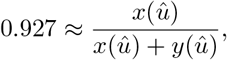

where 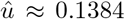 is the blow-up threshold tumor value and *x*(*u*) and *y*(*u*) are computed using the result from Proposition 2 and the post-resection parameter values. Whereas the CT immune cell population remains high in the control case — thereby inhibiting metastatic growth and resulting in a small, dormant metastasis — the perturbation to the system caused by primary tumor resection prompts a short period of transience that includes an increase to the CT immune fraction (from the inflammation) followed by a dip (from tumor-induced immune cell phenotypic plasticity) and a short recovery. For lower levels of inflammation (green curves) the CT immune cell fraction recovers *above* the blow-up threshold, allowing for an effective anti-tumor immune response, capable of inhibiting metastatic blow-up. In contrast, metastatic blow-up occurs whenever the dip in CT immune cell fraction does not recover above the blow-up threshold, 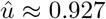. In all cases, the period of transience — and therefore the determination of decay or blow-up — is complete in at most three weeks after the primary resection (Figure 12, inset). Mathematically, this is a consequence of the different time scales: resection is a perturbation of the dynamics from the slow manifold, and the transience is a result of the fast dynamics settling onto the post-resection slow manifold. Biologically, this suggests that the makeup of immune cell populations at potential sites of metastasis can successfully distinguish between metastatic decay and (delayed) blow-up in as little as three weeks after the primary resection.

**Fig. 12.**
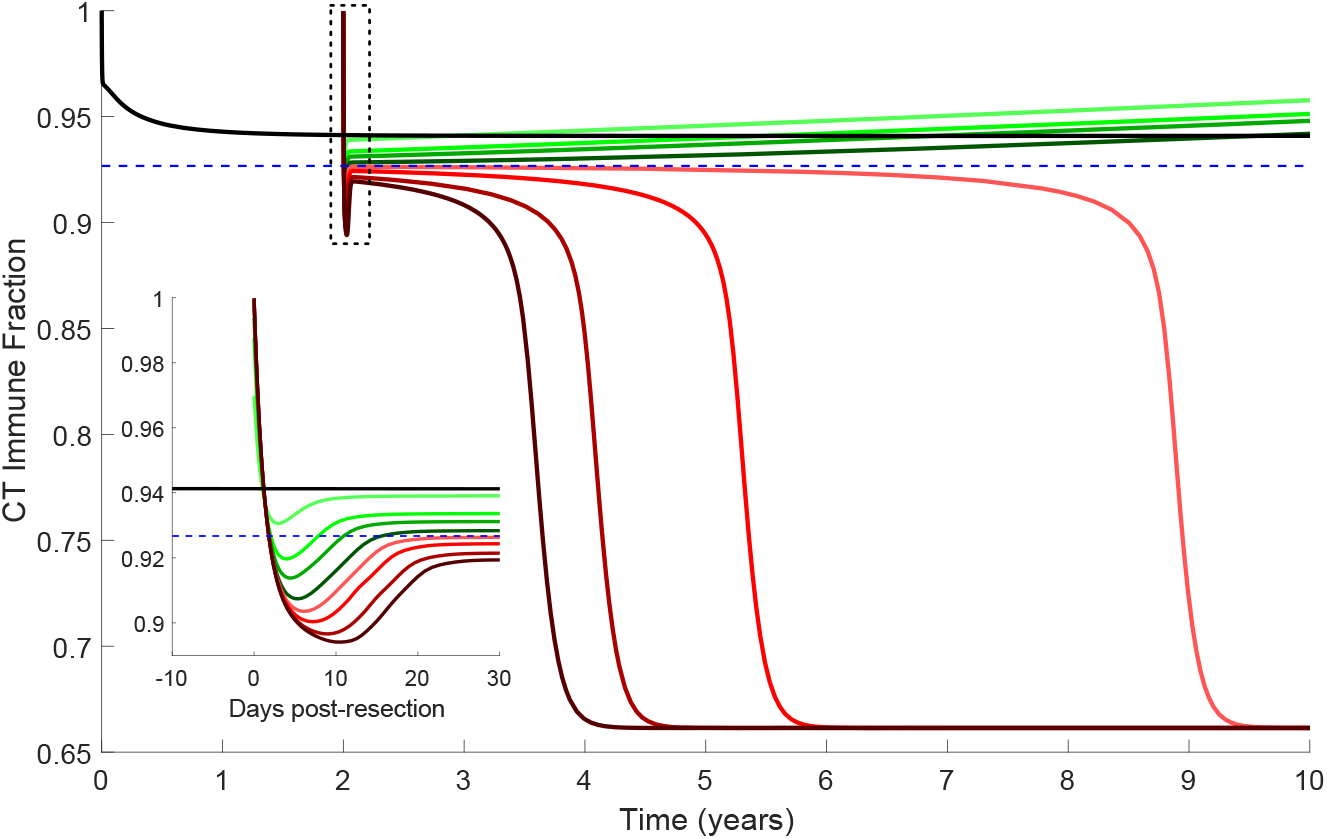
Fraction of CT immune cells at the metastatic site (as a fraction of total immune cell population) as a function of time for the same simulation presented in Figure 10. Horizontal dashed blue line denotes the fraction of CT immune cells predicted at the non-zero unstable node for the post-resection parameter values (92.7% CT immune cells). Inset is a detail of the boxed area of the main figure. Black lines corresponds to control (no resection) dynamics. Colors represent different strengths of the inflammation response to resection and are the same as in Figure 10. Parameters as in Table 1, with the exception of *max*_1_ = 2.5, *r* = 2 × 10^−4^, and *ϕ* = 1.8519 × 10^−5^. Color figure available online.

### 5.4 Combination Treatments and Control of Immune Cells

Diagnosis and prediction are important, but remain toothless without effective treatment strategies. In addition to the strategy of reducing the inflammation associated with primary tumor resection (see Figure 10), the results from the previous section (and our previous work [47]) suggest that targeting the tumor-induced immune cell phenotypic plasticity (tumor education) may be another valid strategy. In the top panel of Figure 13, the time from primary resection to metastatic blow-up (defined here as the metastatic tumor reaching a density of 0.5) is shown as a function of primary resection-induced inflammation (vertical axis) and the rate of tumor education (horizontal axis). Areas in white correspond to parameter values in which blow-up is impossible, or occurs only after 20 years or longer. Times less than 20 years are as indicated. While there are regions where metastatic blow-up is delayed for a decade or longer, the topmost panel of Figure 13 is dominated by delays of 5 years or under, matching the clinical data mentioned in the previous section [46].

**Fig. 13.**
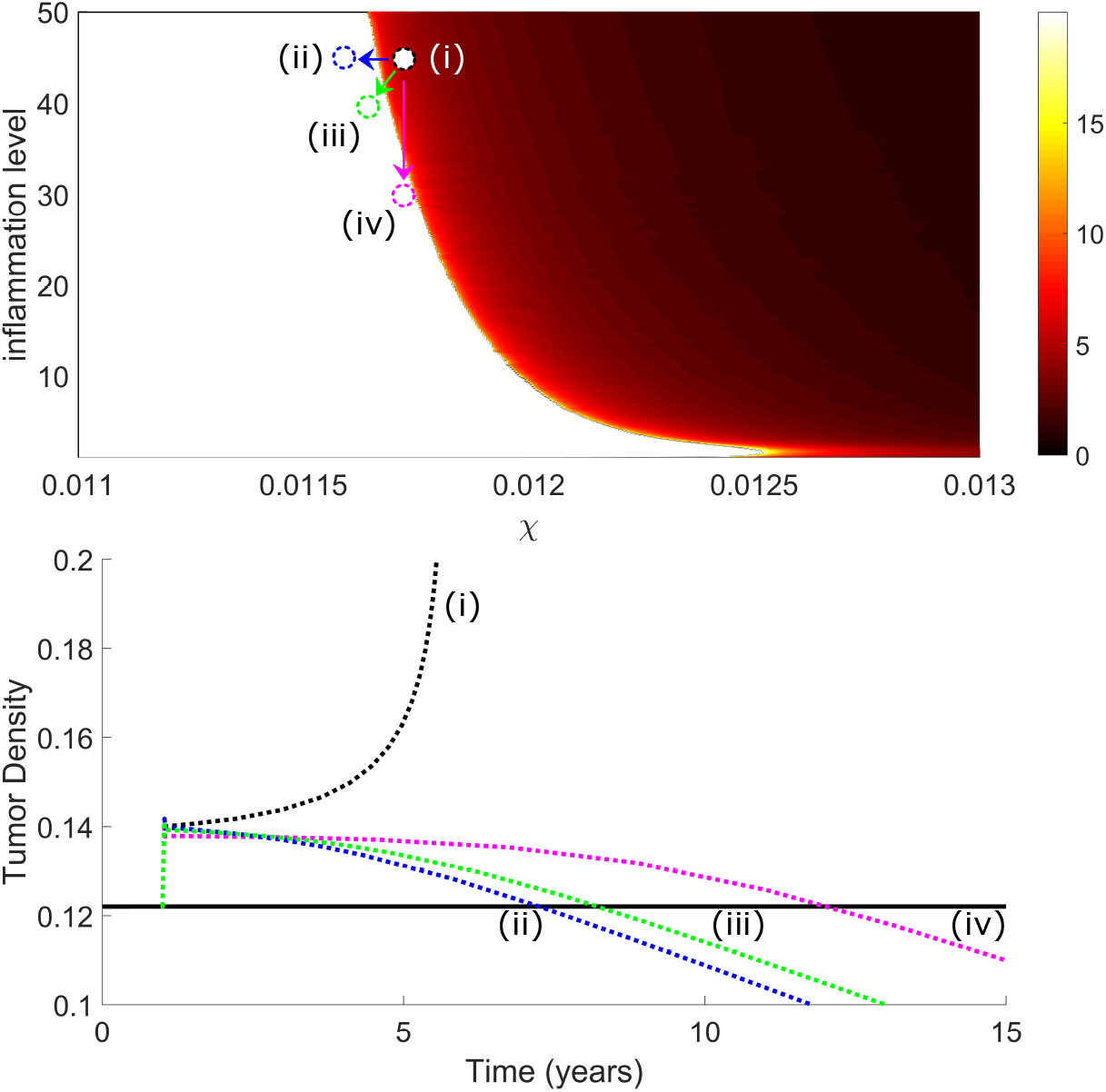
Times (in years post-resection) to metastatic blowup as a function of both the strength of the inflammatory response to primary resection (vertical axis) and the education rate, *χ*, of the CT immune cells by the tumor (horizontal axis). Any times greater than 20 years have been depicted in white. Times defined as the time it takes for tumor density to reach 0.5. The parameters responsible for the four dotted curves in the bottom plot are depicted in the top by circles of corresponding colors. The solid black line in the bottom plot are the control (no resection) dynamics. Dashed black (i) is a reference parameter set in which blow-up is observed. The blue (ii) and magenta (iv) curves are obtained by decreasing only the education rate (to 0.116) or the inflammation (to 30), respectively. The green (iii) curve represents a case in which both the education rate and the inflammation levels are decreased (to 0.1165 and 40, respectively). Control parameters as in Figure12. Color figure available online.

The parameter pair indicated by (i) serves as a reference parameter set for which blow-up is expected, and from which we can evaluate different treatment strategies. In line with experimental [41] and clinical [17,46] evidence, our model predicts that decreasing the inflammation associated with primary tumor resection — moving *down* from (i) to (iv) — can delay blow-up or prevent it entirely. Similar effects can also be achieved by *decreasing* the ability of the tumor to educate CT immune cells — moving *left* from (i) to (ii). By combining these two strategies — (i) to (iii) — similar anti-tumor effects can be achieved using therapeutic strengths that would fail if applied in isolation. Solution trajectories corresponding to the strategies (i)–(iv) are presented in the lower panel of Figure 13.

Although only speculative at this point, our theoretical results concerning the prediction and treatment of metastatic disease *prior* to clinical detection are intriguing and warrant further biological investigation.

## 6 Discussion

As the major cause of cancer-related deaths, a full understanding of the metastatic process is of significant value. Multiple recent lines of investigation — including the discovery of the PMN [29], early metastatic seeding [21,23,48], and decreased metastasis in response to treatments with NSAIDs [28,36,46] — have brought the long-held view of metastasis as the final, unavoidable step in cancer progression into question, and spurred new experimental and theoretical investigations of the process. Two phenomena that are of particular interest are those of metastatic *dormancy* and *blow-up*. Although numerous theories have been proposed to explain these phenomena, and several mathematical models developed to investigate them (notably those of Benzekry and Hanin), prior to the present work, there had not been any mathematical models developed that specifically interrogated the potential roles of pro-tumor immune effects and inflammation.

To fill this lacuna, we have introduced, analyzed, parameterized, and simulated a model of tumor-immune dynamics at the site of a metastatic tumor, informed by the immune-mediated theory of metastasis [47,49]. The model includes both anti- and pro-tumor immune effects, in addition to the experimentally observed phenomena of tumor-induced immune cell phenotypic plasticity — which we have referred to as tumor “education” of anti-tumor immune cells to play a pro-tumor role. By exploiting the difference in timescales between *fast* immune dynamics and *slow* tumor dynamics [23,42], we used techniques from geometric singular perturbation theory and quasi-steady state analysis to reduce the complexity of our model, allowing for meaningful analysis. In particular, we made use of our analytical results to determine conditions necessary for our model to exhibit metastatic dormancy and blow-up.

Within our model, dormancy was possible via two separate mechanisms: either as a small, positive steady state (as in [32]), or as a transient state that occurred as the dynamics traveled near a saddle point (as in [55]). For blow-up, a number of conditions had to be met. First, without any form of primary intervention, the model had to predict a small, dormant metastatic tumor. In the absence of this condition, primary resection could only *hinder* further metastatic growth (Figure 7). Second, blow-up was impossible without assuming a change in the model parameters in response to the primary resection (Figure 9). We tested three possible mechanisms: increased *recruitment* of pro-tumor immune cells, decreased pro-tumor immune cell death rate, and *increased* rate of tumor education of CT immune cells. Only the final strategy was able to successfully reproduce biologically realistic post-resection dormancy and (delayed) blow-up (Figure 10). Third, the primary resection had to induce a short period of growth at the metastatic site, so that a threshold tumor density (unstable node) was passed, and the solution entered the basin of attraction of a larger steady state (Figure 11). In particular, the naive model of primary resection used in previous studies [14,47,51] needed to be replaced with a slightly more sophisticated model. The inclusion of systemic inflammation caused by primary resection surgery (or even a biopsy of the primary tumor), together with tumor “education” of CT immune cells, made it such that metastatic blow-up was possible (Figures 11, and 10).

The claim of Riggi et al. [48] that “the latency period for metastatic tumor development … argues against any form of [pre-metastatic] niche formation” is contested by our results. Figure 12 demonstrates that a significant population of TE immune cells arrives at the site of future metastasis *prior* to significant tumor colonization, thereby creating a permissive PMN. Furthermore, Figure 10 shows that extended metastatic dormancy and subsequent blow-up are still possible, despite (and in fact, largely *because of*) primary tumor removal. While it is true that the removal of the primary tumor would stop any niche preparation that had been underway, our model suggests that the groundwork was already complete by that time. Moreover, it is the inflammation incurred as a result of the resection that jump starts further metastatic growth. In other words, the concepts of PMN and delayed blow-up need not be mutually exclusive, as suggested in [48] (although some of this confusion may be a consequence of loose definitions).

Our results mirror those of both Benzekry [3] and Hanin [23], which demonstrated that an *increase* to metastatic growth after primary resection was needed to explain observed data. In contrast to the results cited, however, our modeling framework allows us to suggest an explicit mechanism for this post-resection increase in metastatic growth and subsequent metastatic blow-up. The initial escape from dormancy is triggered by the inflammation resulting from primary tumor resection. Subsequent growth and eventual metastatic blow-up is sustained by a strong population of pro-tumor TE immune cells at the metastatic site, which is maintained by a post-resection increase to tumor-mediated immune cell phenotypic plasticity. Our results provide support for the theory of metastatic blow-up as a consequence of a systemic inflammatory response to primary tumor resection. There are now mathematical works that have investigated each of the four proposed mechanisms underlying metastatic blow-up described in Section 1, and further work — both experimental and theoretical — must be done to more strongly distinguish between these hypotheses.

Thanks to our sensitivity analysis (Figures 5 and 6), bifurcation analysis (Figure 4), and the accurate reproduction of biologically feasible behavior within reasonable parameter constraints, we posit that the biological predictions discussed herein are biologically realistic and of significant potential value. In particular, the concept of a post-resection change in metastatic behavior has now been demonstrated in at least three distinct theoretical settings (this work, [3], and [23]), and warrants further targeted biological investigation. The observations of den Breen and Eftimie [5] that the ratio of pro-tumor to anti-tumor macrophages may be a valuable predictive measure were also found herein, this time in the metastatic setting. Indeed, within three weeks of the primary tumor resection, and well before the metastatic tumor is clinically detectable, the immune cell composition at a potential site of metastasis can predict whether or not metastatic blow-up will occur upwards of a decade before the tumor would be clinically detectable (Figure 10). Although compelling, further biological studies are needed in order to confirm our theoretical results, including the determination of (i) whether a post-resection jump in pro-tumor immune cells at distant sites is actually observed, and, (ii) if such a jump does occur, what are the mechanisms responsible?

In addition to confirming previous experimental [41] and clinical [46] results by reproducing the anti-metastasis effects of inhibiting inflammation associated with primary tumor resection (Figures 10 and 13), our model also provides another potential target for therapy: inhibiting tumor-induced immune cell phenotypic plasticity [39] (Figure 13). While intriguing, this result also does have its limitations. For instance, what exactly is meant by “tumor-induced immune cell phenotypic plasticity,” what are the mechanisms underlying it, and is it reversible? Answers to these questions must first be found before putative therapeutic targets can be suggested.

Whereas the results presented in this work are exciting, we stress that caution should be taken when interpreting the results as they are based on a *model* which relies on many simplifications and assumptions. First and foremost, we have used a non-spatial ODE model to describe an inherently spatial process. While this was done in order to simplify the model to allow for deeper analysis, future work should certainly include the effects of space as well as the architecture of the body. Such considerations have been taken previously, with a spatially explicit stochastic model introduced by Frei et al. [20], a detailed multi-site ODE model of tumor-immune dynamics developed by Poleszczuk, Walker, and colleagues [43,44,52,51] and, combining the approaches, Franßen et al. [19,18] have suggested a multi-site PDE model. Future work includes adding pro-tumor immune effects in such modeling frameworks.

## Acknowledgements

AR gratefully acknowledges support from an Alberta Innovates Graduate Student Scholarship. TH is supported by an NSERC Discovery grant.

## A Appendix

Here we prove Proposition 3.

*Proof* Consider 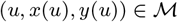. We compute the derivatives *x_u_*(*u*) and *y_u_*(*u*). We begin with *x_u_*:

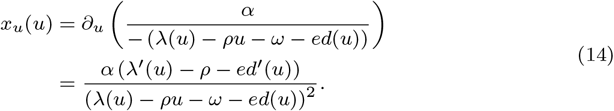

The sign of *x_u_* depends entirely on the sign of

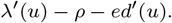

With the choices we made for *λ*(*u*) and *ed*(*u*), we arrive at the following chain of equivalent conditions:

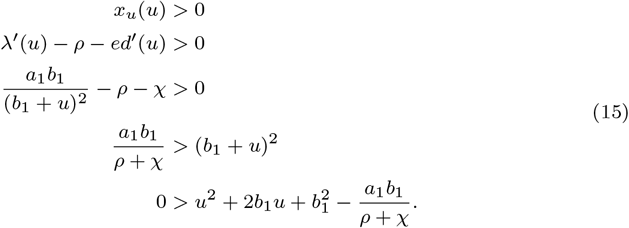

The roots to the above quadratic are given by

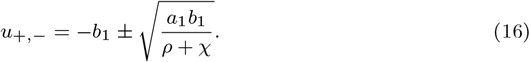

This gives distinct, real roots (assuming that *a*_1_*b*_1_ ≠ 0) with at least one of them negative. If *u*_+_ > 0, then we see that *x_u_* > 0 for *u < u*_+_ and negative otherwise. If *u*_+_ < 0, then we simply have *x_u_* < 0 for all non-negative *u*.

Next, we consider *y_u_*. We show simply that *y_u_* is non-negative, as

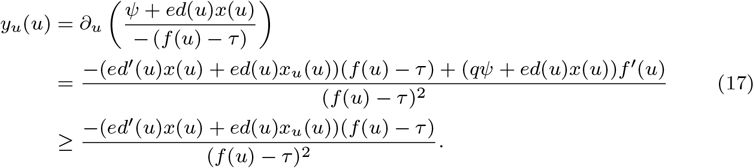

The final inequality results from the fact that (*ψ* + *ed*(*u*)*x*(*u*))*f*′(*u*) ≥ 0 since *f* is increasing (A1). Now, we can use the expressions for *x*(*u*) and *x_u_*, as well as our choice of *ed*(*u*) = *χ^u^* to arrive at

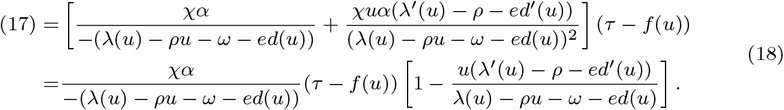

The sign of this expression depends only on the sign of the term

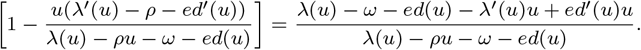

By assumption (A2), the denominator is always negative, and therefore the sign of the above expression is determined by the sign of its numerator. With the choices for *ed*(*u*) and *λ*(*u*) from Section 2.3, the numerator simplifies to

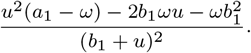

Using the fact that *a*_1_ − *ω* < 0 (A2) guarantees that the quadratic is negative for all *u* ≥ 0, and therefore *y_u_*(*u*) ≥ 0 for all *u* ≥ 0.

## References

1. Balkwill, F., Mantovani, A.: Inflammation and cancer: back to Virchow? Lancet 357, 539–545 (2001)

2. Benzekry, S., Gandolfi, A., Hahnfeldt, P.: Global dormancy of metastases due to systemic inhibition of angiogenesis. PLoS One 9(1), e84249 (2014)

3. Benzekry, S., Lamont, C., Barbolosi, D., et al.: Mathematical modeling of tumor-tumor distant interactions supports a systemic control of tumor growth. Cancer Research 77(18), 5183–5193 (2017)

4. Blomberg, O., Spagnuolo, L., de Visser, K.: Immune regulation of metastasis: mechanistic insights and therapeutic opportunities. Disease Models and Mechanisms (2018)

5. den Breems, N., Eftimie, R.: The re-polarisation of m2 and m1 macrophages and its role on cancer outomes. Journal of Theoretical Biology 390, 23–39 (2016). Doi:10.1016/j.jtbi.2015.10.063

6. Cameron, M., et al.: Temporal progression of metastasis in lung: Cell survival, dormancy, and location dependence of metastatic inefficiency. Cancer Research 60(9), 2541–2546 (2000)

7. Chambers, A., Groom, A., MacDonald, I.: Dissemination and growth of cancer cells in metastatic sites. Nature Reviews Cancer 2(8), 563–572 (2002)

8. Chiarella, P., Bruzzo, J., Meiss, R., Ruggiero, R.: Concomitant tumor resistance. Cancer Letters 324(2), 133–141 (2012)

9. Cohen, E., Gao, H., Anfossi, S., et al.: Inflammation mediated metastasis: Immune induced epithelial-to-mesenchymal transition in inflammatory breast cancer cells. PLoS One (e0132710) (2015)

10. Colegio, O., Chu, N., Szabo, A., et al.: Functional polarization of tumor-associated macrophages by tumour-derived lactic acid. Nature 513, 559–563 (2014)

11. Coughlin, C., Murray, J.: Current and emerging concepts in tumour metastasis. The Journal of Pathology 222(1) (2010). Doi:10.1002/path.2727

12. Del Monte, U.: Does the cell number 10^9^ still really fit one gram of tumor tissue? Cell Cycle 8(3), 505–506 (2009). Doi:

13. Dewan, M., Galloway, A., Kawashima, N., et al.: Fractionated but not single-dose radiotherapy induces an immune-mediated abscopal effect when combined with anti-ctla-4 antibody. Clinical Cancer Research 15(17), 5379–5388 (2009). Doi:10.1158/1078-0432.CCR-09-0265

14. Eikenberry, S., Thalhauser, C., Kuang, Y.: Tumor-immune interaction, surgical treatment, and cancer recurrence in a mathematical model of melanoma. PLoS Computational Biology 5(4), e1000362 (2009). Doi:10.1371/journal.pcbi.1000362

15. El-Kenawi, A., Gatenby, C., Robertson-Tessi, M., et al.: Acidity promotes tumour progression by altering macrophage phenotype in prostate cancer. British Journal of Cancer (2019). DOI: 10.1038/s41416-019-0542-2

16. Emens, L., Ascierto, P., Darcy, P., et al.: Cancer immunotherapy: Opportunities and challenges in the rapidly evolving clinical landscape. European Journal of Cancer 81, 116–129 (2017)

17. Forget, P., Vandengende, J., Berliere, M., et al.: Do intraoperative analgesics influence breast cancer recurrence after mastectomy? a retrospective analysis. Anesthesia and Analgesia 110(6), 1630–1635 (2010)

18. Franßen, L., Chaplain, M.: A mathematical multi-organ model for bidirectional epithelial-mesenchymal transitions in the metastatic spread of cancer. bioRxiv (2019). DOI: 10.1101/745547

19. Franßen, L., Lorenzi, T., Burgess, A., et al.: A mathematical framework for modelling the metastatic spread of cancer. Bulletin of Mathematical Biology (81), 1965–2010 (2019)

20. Frei, C., Hillen, T., Rhodes, A.: A stochastic model for cancer metastasis: Branching stochastic process with settlement. Mathematical Medicine and Biology (2018). Accepted. BioRXiv: 10.1101/294157

21. Friberg, S., Nystrom, A.: Cancer Metastases: Early Dissemination and Late Recurrences. Cancer Growth and Metastasis (8), 43–49 (2015)

22. Gorelik, E.: Concomitant tumor immunity and the resistance to a second tumor challenge. Advances in Cancer Research 39, 75–120 (1983)

23. Hanin, L., Rose, J.: Suppression of metastasis by primary tumor and acceleration of metastasis following primary tumor resection: A natural law? Bulletin of Mathematical Biology 80(3), 519–539 (2018)

24. Hanin, L., Seidel, K., Stoevesandt, D.: A universal model of metastatic cancer, its parametric forms and their identification: what can be learned from site-specific volumes of metastases. Journal of Mathematical Biology 72, 1633–1662 (2016). Doi:10.1007/s00285-015-0928-6

25. Hek, G.: Geometric singular perturbation theory in biological practice. Journal of Mathematical Biology 60(3), 347–386 (2010)

26. Hobson, J., Gummadidala, P., Silverstrim, B., et al.: Acute inflammation induced by the biopsy of mouse mammary tumors promotes the development of metastasis. Breast Cancer Research and Treatment 139(2), 391–401 (2013)

27. Jones, C.: Geometric singular perturbation theory. In: J. Russell (ed.) Dynamical Systems, pp. 44–118. 2nd Session of the Centro Internazionale Matematico Estivo (CIME), Berlin: Springer, Montecatini Terme, Italy (1995)

28. Joyce, J., Pollard, J.: Microenvironmental regulation of metastasis. Nat. Rev. Cancer 9(4), 239–252 (2009). Doi:10.1038/nrc2618

29. Kaplan, R., Riba, R., Zacharoulis, S., et al.: VEGFR1-positive haematopoietic bone marrow progenitors initiate the pre-metastatic niche. Nature 438, 820–827 (2005). Doi:10.1038/nature04186

30. Kim, Y., Jeon, H., Othmer, H.: The role of the tumor microenvironment in glioblastoma: A mathematical model. IEEE Transactions on Bio-Medical Engineering 64(3), 519–527 (2017)

31. Kumar, G., Manjunatha, B.: Metastatic tumors to the jaws and oral cavity. Journal of Oral and Maxillofacial Pathology 17(1), 71–75 (2013)

32. Kuznetsov, V., Makalkin, V., Taylor, I., et al.: Nonlinear dynamics of immunogenic tumors: Parameter estimation and global bifurcation analysis. Bulletin of Mathematical Biology 50(2), 295–321 (1994)

33. Liu, V., Wong, L., Jang, T., et al.: Tumor Evasion of the Immune System by Converting CD4+CD25-T Cells into CD4+CD25+ T Regulatory Cells: Role of Tumor-Derived TGF-*β*. The Journal of Immunology 178, 2883–2892 (2007)

34. Luzzi, K., et al.: Multistep nature of metastatic inefficiency: Dormancy of solitary cells after successful extravasation and limited survival of early micrometastases. American Journal of Pathology 153(865) (1998). DOI: 10.1016/S0002-9440(10)65628-3

35. Maida, V., Ennis, M., Kuziemsky, C., Corban, J.: Wounds and survival in cancer patients. European Journal of Cancer 45(18), 3237–3244 (2009)

36. Marx, J.: Inflammation and cancer: The link grows stronger. Science 306(5698), 966–968 (2004)

37. de Mingo Pulido, A., Ruffell, B.: Immune regulation of the metastatic process: Implications for therapy. Advances in Cancer Research 132, 139–163 (2016)

38. Norton, K., Jin, K., Popel, A.S.: Modeling triple-negative breast cancer heterogeneity: Effects of stromal macrophages, fibroblasts and tumor vasculature. Journal of Theoretical Biology 452, 56–68 (2018)

39. Oleinika, K., Nibbs, R., Graham, G., et al.: Suppression, subversion and escape: The role of regulatory t cells in cancer progression. Clinical and Experimental Immunology 171(1), 36–45 (2013)

40. Olobatuyi, O., de Vries, G., Hillen, T.: A reaction-diffusion model for radiation-induced bystander effects. Journal of Mathematical Biology 75(2), 341–372 (2017)

41. Park, C., Hartl, C., Schmid, D., et al.: Extended release of perioperative immunotherapy prevents tumor recurrence and eliminates metastases. Science Translational Medicine 10(433), eaar1916 (2018)

42. Pascual, M., Alonso, S., Pares, D., et al.: Randomized clinical trial comparing inflammatory and angiogenic response after open versus laparoscopic curative resection for colonic cancer. British Journal of Surgery 98, 50–59 (2010)

43. Poleszczuk, J., Luddy, K., Prokopiou, S., et al.: Abscopal benefits of localized radiotherapy depend on activated t-cell trafficking and distribution between metastatic lesions. Cancer Research 76(5), 1009–1018 (2016). Doi: 10.1158/0008-5472.CAN-15-1423

44. Poleszczuk, J., Moros, E., Fishman, M., et al.: Modeling T-cell trafficking to increase the likelihood of radiation-induced abscopal effects. Journal of Targeted Therapies in Cancer 06.17, 36–40 (2017)

45. Prehn, R.: Two competing influences that may explain concomitant tumor resistance. Cancer Research 53(14), 3266–3269 (1993)

46. Retsky, M., Demicheli, R., Hrushesky, W., et al.: Reduction of breast cancer relapses with perioperative non-steroidal anti-inflammatory drugs: new findings and a review. Current Medicinal Chemistry 20(33), 4163–4176 (2013)

47. Rhodes, A., Hillen, T.: A mathematical model of the immune-mediated theory of metastasis. Journal of Theoretical Biology (2019). Submitted. Pre-print available at bioRxiv, DOI: 10.1101/565531

48. Riggi, N., Auget, M., Stamenkovic, I.: Cancer metastasis: A reappraisal of its underlying mechanisms and their relevance to treatment. Annual Review of Pathology: Mechanisms of Disease 13, 117–140 (2018)

49. Shahriyari, L.: A new hypothesis: some metastases are the result of inflammatory processes by adapted cells, especially adapted immune cells at sites of inflammation. F1000 Research 5(175) (2016). Doi:10.12388/f1000research.8055.1

50. Tyzzer, E.: Factors in the production and growth of tumor metastases. Journal of Medical Research 28(2), 309–332 (1913)

51. Walker, R., Poleszczuk, J., Pilon-Thomas, S., et al.: Immune interconnectivity of anatomically distant tumors as a potential mediator of systemic responses to local therapy. Scientific Reports 8(9474) (2018)

52. Walker, R., Schoenfeld, J., Pilon-Thomas, S., et al.: Evaluating the potential for maximized t cell redistribution entropy to improve abscopal responses to radiotherapy. Convergent Science Physical Oncology 3(034001) (2017)

53. Walter, N., Rice, P., Redente, E., et al.: Wound healing after trauma may predispose to lung cancer metastasis: Review of potential mechanisms. American Journal of Respiratory Cell and Molecular Biology 44(5), 591–596 (2011). Doi: 10.1165/rcmb.2010-0187RT

54. Weiss, L.: Metastatic inefficiency. Advances in Cancer Research 54, 159–211 (1990)

55. Wilkie, K., Hahnfeldt, P.: Modeling the dichotomy of the immune response to cancer: Cytotoxic effects and tumor-promoting inflammation. Bulletin of Mathematical Biology 79(6), 1426–1448 (2017)

56. Yang, Z., Tang, X., Meng, G., et al.: Latent cytomegalovirus infection in female mice increases breast cancer metastasis. Cancers 11(447), 1–20 (2019)

